# Transforming growth factor-beta signaling via ALK1 and ALK5 regulates distinct functional pathways in vein graft intimal hyperplasia

**DOI:** 10.1101/860320

**Authors:** EL Low, JT Schwartze, A Kurkiewicz, M Pek, DJ Kelly, AS Shaw, M Thorikay, J McClure, M McBride, S Arias-Rivas, SE Francis, NW Morrell, C Delles, P Herzyk, MJ Havenga, SA Nicklin, P Ten Dijke, AH Baker, AC Bradshaw

## Abstract

**Rationale:** Transforming growth factor-beta (TGFβ) is tightly regulated at multiple levels, with regulation at the receptor level now recognized as a key determinant of the cellular response to this pleiotropic cytokine. TGFβ promotes saphenous vein graft neointima formation after coronary artery bypass graft (CABG) surgery, inducing smooth muscle cell (SMC) hyperplasia and fibrosis by signaling via activin receptor-like kinase 5(ALK5). However, the role of the alternate TGFβ receptor ALK1 remains completely unknown.

**Objective:** To define the receptor pathways activated by TGFβ in SMCs and their mechanistic importance during CABG neointima formation.

**Methods and results:** Radioligand co-IP assays revealed direct interactions between TGFβ, ALK5 and ALK1 in primary saphenous vein graft SMC (HSVSMC) from patients undergoing CABG. Knockdown and pharmacological inhibition of ALK5 or ALK1 in HSVSMC significantly attenuated TGFβ-induced phosphorylation of receptor-regulated (R)-Smads 2/3 and 1/5, respectively. Microarray profiling followed by qRT-PCR validation showed that TGFβ induced distinct transcriptional networks downstream of ALK5 or ALK1, associated with HSVSMC contractility and migration, respectively and confirmed using migration assays as well as qRT-PCR and western blot assays of contractile SMC markers. scRNAseq analysis of TGFβ-treated HSVSMC identified distinct subgroups of cells showing ALK5 or ALK1 transcriptional responses, while RNA velocity analyses indicated divergence in differentiation towards ALK5 or ALK1-dominant lineages. ALK1, ALK5 and their downstream effectors pSmad1/5 and pSmad2/3 were localized to αSMA+ neointimal SMCs in remodelled mouse vein grafts. Pharmacological inhibition or genetic ablation of Smad1/5 substantially reducing neointima formation following acute vascular injury. Notably, expression and activation of ALK1, ALK5 and their respective downstream R-Smads was already evident in hyperplastic saphenous veins prior to grafting.

**Conclusions:** Whilst canonical TGFβ signaling via ALK5 promotes a contractile HSVSMC phenotype, transactivation of ALK1 by TGFβ induces neointima formation by driving cell migration. Restoring the balance between ALK1 and ALK5 in HSVSMC may represent a novel therapeutic strategy for vein graft failure.

## Introduction

Coronary artery bypass grafting (CABG) is a mainstay in the treatment of coronary artery disease (CAD), with over 400,000 CABG surgeries performed per annum in the United States alone [1]. However, fewer than 60% of saphenous vein grafts (SVG) remain patent in the long-term (>10yrs) due to the formation of a hyperplastic, occlusive neointima within the grafted vessel [2]. Smooth muscle cell (SMC) hyperplasia and matrix biosynthesis are key drivers of vein graft intimal hyperplasia (IH), which is a fertile substrate for the accelerated development of rupture-prone atherosclerotic plaques [3, 4]. SMC hyperplasia has therefore been identified as an important target for therapeutic interventions aimed at improving outcomes after CABG surgery [5].

The pleiotropic cytokine transforming growth factor-beta (TGFβ1) is highly expressed in re-stenotic vessels from CAD patients and is acutely upregulated following vein graft implantation in animal models of vein graft disease, inducing constrictive and fibrotic vascular remodelling [6–8]. However, to date most studies investigating the role of TGFβ in IH have used global approaches to target TGFβ, or have focused on the canonical TGFβ type I receptor (also termed activin receptor-like kinase 5/ALK5), Smad2/3-mediated pathway. TGFβ elicits a diverse range of cellular responses by activating different signaling pathways, and in certain cell types can also signal via a distinct type I receptor, activin receptor-like kinase 1(ALK1), activating a separate set of receptor-regulated (R)-Smad proteins (Smad1/5) that can antagonize ALK5 signaling [9]. Interestingly, a recent study showed that vascular injury-induced upregulation of KLF6 (Kruppel-like growth factor 6) in endothelial cells (ECs) was associated with increased expression of ALK1 in neighboring SMC [10]. Moreover, mutations in exon 10 of the *ACVRL1* gene (encodes ALK1) are associated with pulmonary arterial hypertension, a fatal condition that is characterized by the development of IH within the small pulmonary arteries [11]. Although this points to a potentially relevant role for ALK1 in SMC during neointima formation, studies have so far failed to show whether TGFβ is able to signal via ALK1 in human SMCs. The role of SMC ALK1 during the development of vein graft IH has also yet to be defined. Here we show that whilst TGFβ signaling via ALK5 is associated with SMC contractility and fibrosis, activation of ALK1 promotes neointima formation by inducing a distinct transcriptional network that drives SMC migration, identifying this pathway as a potential therapeutic target in the setting of vein graft failure (VGF).

## Results

### TGFβ1 signals through ALK1 in saphenous vein SMCs

The expression pattern and abundance of TGFβ receptors varies substantially between cell types, influencing the cellular response to TGFβ stimulation. We therefore set out to characterize the expression and activation of TGFβ receptors in SMC outgrowth cultures (HSVSMC) from saphenous vein segments obtained from patients undergoing CABG surgery. *ALK1, ALK5*, *TβR2* (TGFβ type II receptor) and the accessory TGFβ receptors endoglin (*ENG*) and betaglycan (*TβR3*) were widely expressed in HSVSMC at the RNA level (n=7-12 patients, Fig.1A). Affinity labeling and crosslinking studies using I^125^TGFβ indicated binding of TGFβ to all these receptors in HSVSMC, with particularly strong binding to ALK5, ENG and TβRIII/ TβR3 (Fig.1B). Activation of ALK receptors induces R-Smad phosphorylation, initiating a cascade of receptor-specific downstream events that culminate in the translocation of phosphorylated R-Smad proteins to the nucleus [12]. Immunoblotting revealed that treatment with even low doses of TGFβ (0.1ng/mL) could induce a rapid increase in pSmad1/5 as well as pSmad2 (Fig.1C, upper [pSmad2] and lower [pSmad1/5] panels), while immunocytochemistry confirmed the nuclear localization of both pSmad1/5 and pSmad2/3 signaling complexes following stimulation of HSVSMC with TGFβ (Fig.1D). QRT-PCR evaluation of the prototypical ALK1 and ALK5 gene targets inhibitor of differentiation-1(*ID1*; ALK1/pSmad1/5 target) and plasminogen activator inhibitor-1(*PAI1*/*SERPINE1*; ALK5/pSmad2/3 target) was consistent with activation of both pathways by TGFβ at the transcriptional level (Fig.1E and 1F). Importantly, pharmacological inhibition (using the ALK1 kinase inhibitor KO2288/KO, or the ALK5 kinase inhibitor SB525334/SB; Supplemental Fig.1A) or dsiRNA-mediated knockdown of ALK1 or ALK5 (Supplemental Fig.1B) in TGFβ-stimulated HSVSMC reduced the expression of pSmad1/5 and pSmad2, respectively, confirming that TGFβ can bind to both ALK1 and ALK5 in HSVSMC, activating separate Smad-mediated signaling pathways.

**Figure 1.**
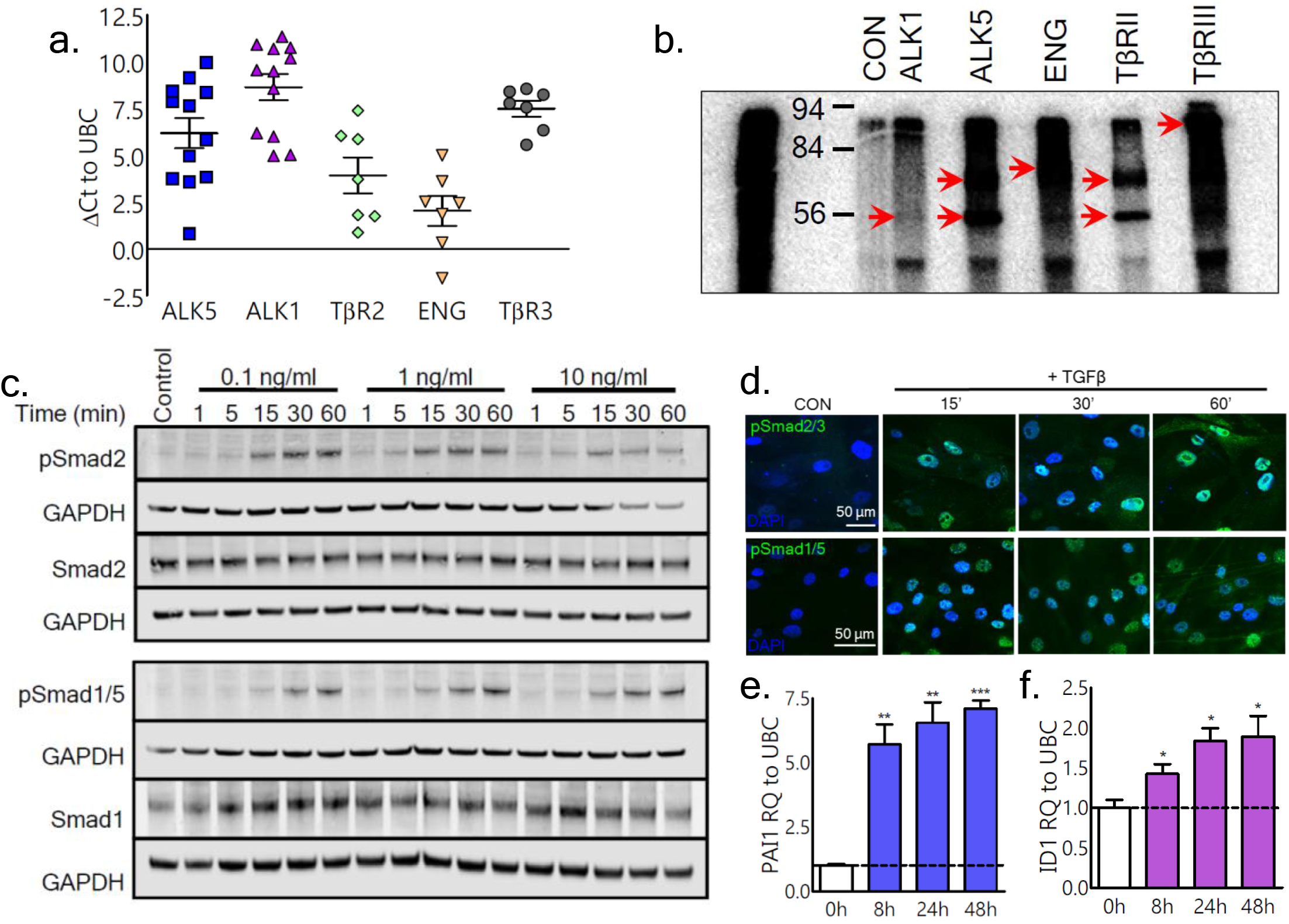
TGFβ signals via both ALK5 and ALK1 pathways in human saphenous vein SMCs. mRNA expression of TGFβ receptors in HSVSMCs from CABG patients was assessed via qRT-PCR (**A**) (data expressed as ΔCt relative to UBC housekeeping gene; n=7-12) HSVSMCs were treated with I^125^ TGFβ prior to biochemical crosslinking and harvesting for immunoprecipitation with TGFβ receptor antibodies as indicated **(B).** Serum-starved HSVSMCs were stimulated with TGFβ for up to 1h for Western blot **(C)** or ICC **(D)** analysis of phosphorylated and unphosphorylated Smad2 or Smad1 (Representative western blot and confocal microscopy ICC images shown; n=3). (**E, F**) qRT-PCR analysis of known gene targets of the ALK5 pathway (PAI1:**E**) and ALK1 pathway (ID1:**F**) were performed following 24h stimulation with TGFβ (n=3; repeated measures one-way ANOVA with Tukeys post-hoc tests, *=p<0.05, **=p<0.001,***= p<0.001 compared to control).

### TGFβ1 regulates distinct subsets of pro-migratory or contractile genes by signaling via ALK1 and ALK5

Having shown that stimulation of HSVSMC with TGFβ induces the rapid activation of both ALK1 and ALK5 pathways, we next aimed to evaluate the effect of ALK1 and ALK5 signaling at the transcriptional level, by performing microarray analysis of HSVSMC stimulated with TGFβ in the presence or absence of pharmacological ALK1 or ALK5 inhibitors (KO2288/KO or SB525334/SB respectively, n=3 patients, venn diagram of 5% FDR [false discovery rate] significant probesets shown in Supplemental Fig.2). Interestingly, Ingenuity Pathway Analysis (IPA) of Limma (linear models for microarray data) significant probes (p<0.05) showed that there was little overlap in the molecular and cellular functions and top canonical pathways regulated by TGFβ signaling via ALK5 or ALK1 (Fig.2A and 2C respectively). Top downregulated genes following ALK5 inhibition included many key mediators of fibrosis, including connective tissue growth factor (*CTGF*) and fibroblast growth factor-2(*FGF2*) (Supplemental Fig.3A and B; qRT-PCR validation of *CTGF*, *FGF2* and other top ALK5 targets shown in Fig.2B), in line with previous studies indicating that ALK5 is responsible for driving the pro-fibrotic actions of TGFβ [13]. Importantly, several genes associated with the contractile SMC phenotype were also highlighted in our array analyses, and qRT-PCR validation in HSVSMC from a further three CABG patients confirmed the ALK5-specific regulation of alpha-smooth muscle actin (*ACTA2*), calponin (*CNN1*), transgelin (*TAGLN*) and smooth muscle myosin heavy chain 11 (*MYH11*) (Figure 2E). S20 myosin light chain subunit phosphorylation (pS20-MLC), a key regulatory event driving SMC contraction, was also reduced following ALK5 inhibitor treatment in TGFβ-stimulated HSVSMCs (Fig.2F). To corroborate these findings, we employed an established contractile SMC differentiation protocol [14]. Western immunoblot analysis confirmed a substantial induction of ACTA2, CNN1 and transgelin (SM22α) (Supplemental Fig.4A), pS20-MLC and total MLC (tMLC; Supplemental Fig.4B) in HSVSMC cultured in smooth muscle differentiation medium (SMDS) compared to 15% FCS SMC growth medium, peaking at the 6 day timepoint. Pharmacological inhibition of ALK5, but not ALK1, significantly downregulated the mRNA expression of *ACTA2* and *CNN1* in SMDS-treated cells at day 6 (p<0.01, Supplemental Fig.4C and D), further confirming the importance of the ALK5 receptor pathway in the regulation of SMC contractility.

**Figure 2.**
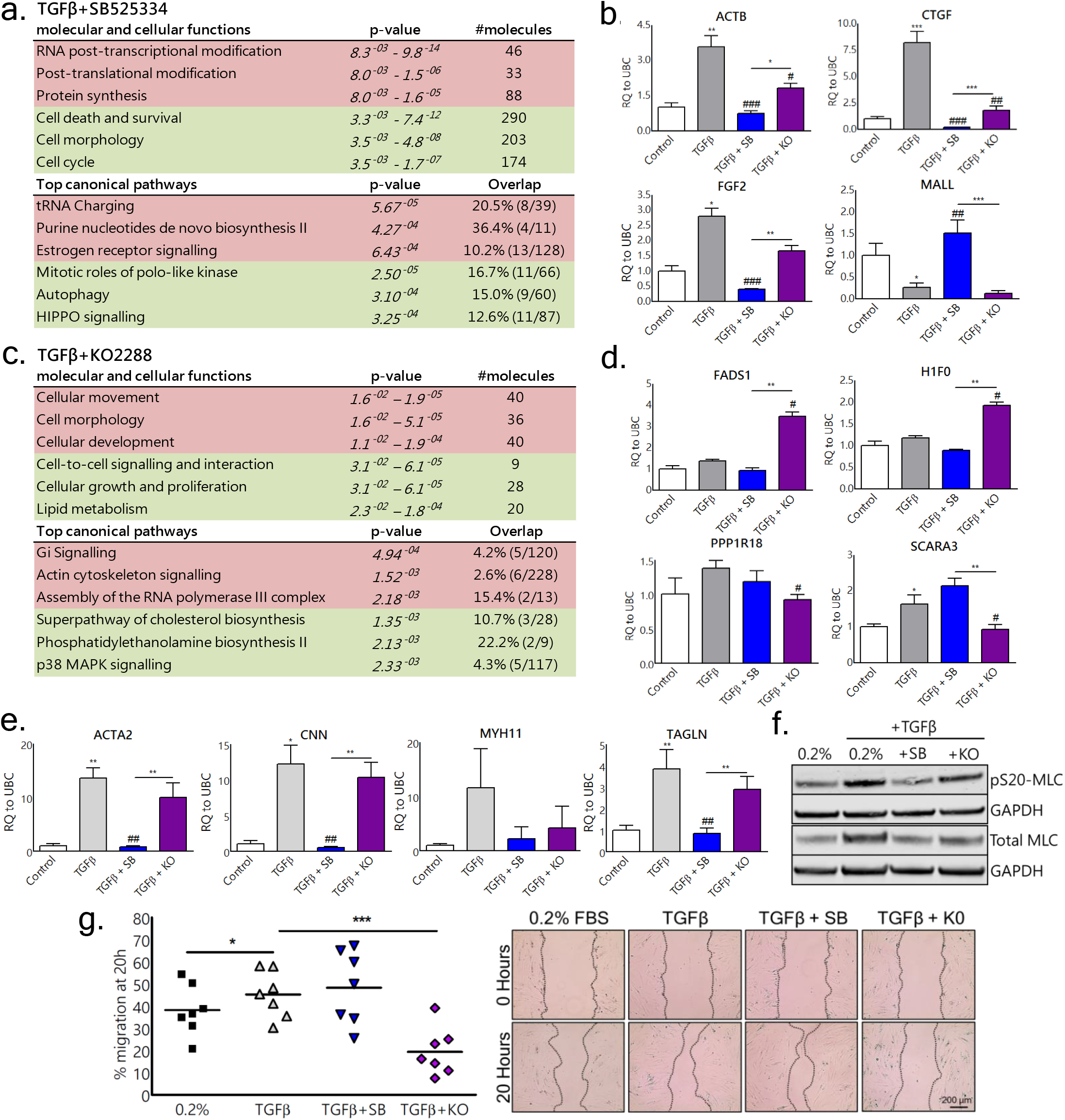
Activation of ALK5 and ALK1 pathways by TGFβ in HSVSMCs induces pathway-specific transcriptional changes associated with distinct functional outcomes. Whole-genome expression profiling using the Illumina HT-12 v4 Expression BeadChip was performed on mRNA extracted from serum-starved HSVSMC stimulated with TGFβ ± ALK5 or ALK1 kinase inhibitors (**SB525334/SB**or **K02288/KO**, respectively) for 24h (n=3). Ingenuity Pathway Analysis identified significantly downregulated (red) and upregulated (green) molecular and cellular functions/canonical pathways based on limma significant probes using a 5 % FDR cut-off **(A, C)**Microarray validation by qRT-PCR was performed on a further 3 patients for top dysregulated genes following ALK5 (ACTB, FGF2, CTGF, MALL; **B**) or ALK1 inhibition (FADS1, H1F0, PPP1R18 and SCARA3; **D**) as well as evaluation of the contractile SMC markers ACTA2, CNN, MYH11 and TAGLN (**E**) (*/#=p<0.05, **/##=p<0.01, ***/###=p<0.001 compared to control(*) or TGFβ (#)). Expression of phosphorylated and total myosin light chain was evaluated by western immunoblotting (representative images from n=3 patients)(**F**). Migration 20h after scratch was compared to 0h timepoint in HSVSMC treated with TGFβ ± inhibitors (*=p<0.05, ***=p<0.001; n=7)(**F**). Data in graphs represent mean ± SEM, analysed by repeated measures one-way ANOVA with Tukeys post-hoc tests.

Top genes identified in our microarray analysis as dysregulated following ALK1 inhibition were also confirmed by qRT-PCR validation (Fig.2D, p<0.05 [*FADS1, H1F0, PPP1R18* and *SCARA3*]) and included regulators of cholesterol transport and biosynthesis (Supplemental Fig.3; tables C and D). Intriguingly, cholesterol biosynthesis and lipid metabolism were highlighted as two functional pathways regulated by TGFβ via ALK1, suggesting that ALK1 in SMC may play a similar role in promoting lipid uptake as recently observed in ECs [15]. Importantly, IPA analysis highlighted ‘cellular movement’ as the top cellular function targeted at the transcriptional level following ALK1 inhibition (Fig.2C, top table). In agreement with this observation, pharmacological inhibition of ALK1 in wound healing assays significantly attenuated TGFβ-mediated HSVSMC migration (Fig.2F, KO2288/KO group; p<0.01, n=7 patients) whereas inhibition of ALK5 led to a slight increase in migration (Fig.2F, SB525334/SB group; p=0.11). Taken together, these data indicate that TGFβ activates two divergent receptor signaling pathways in HSVSMC, inducing migration by activating a specific sub-set of target genes downstream of ALK1 whilst promoting SMC fibrosis and contractility by signaling via ALK5.

### scRNA-seq analysis identifies nonoverlapping ALK5- and ALK1-dominant HSVSMC subgroups

Having identified divergent receptor pathways in HSVSMCs using bulk cell approaches including western blotting, microarray and qPCR analysis, we next aimed to evaluate the heterogeneity of HSVSMC transcriptional responses to TGFβ stimulation using scRNA-seq. Quiesced HSVSMC were stimulated with TGFβ1 for 24 hours prior to harvesting for scRNA-seq using the droplet-based 10x Chromium platform. 5,245 untreated cells and 7,073 TGFβ-treated cells from the same patient were analyzed, with approximately 3000 genes detected per cell in both samples (Supplemental Fig.5). In line with our previous observations, TGFβ-treated cells showed substantially higher expression of well-established TGFβ response genes including *SERPINE1* and *CTGF* (connective tissue growth factor), as well as ALK1 transcriptional targets such as *ID3* (inhibitor of differentiation 3) and *PLAUR* (plasminogen activator) (Fig.3A, L-R, gene names indicated above plots) (Nurgazieva et al, 2014). We observed an uneven distribution of *SERPINE1* and *PLAUR* expression in particular. To further evaluate this heterogeneous expression pattern, cells were projected into 2-dimensional space and unsupervised clustering using the Louvain algorithm was performed, identifying 11 sub-groupings within the TGFβ-treated group (sample 2; Fig.3B; 30 most differentially expressed genes in each subgroup shown in Supplemental Fig. 5). Between these sub-groups, *SERPINE1* was most highly expressed in subgroup 4; conversely, this subgroup showed the lowest expression of *PLAUR* (Fig.3C). Instead, *PLAUR* showed high expression in subgroup 0, which was amongst the lowest-expressing subgroups for *SERPINE1* expression and located at the opposite end of the UMAP Louvain plot (Fig.3C, upper and lower panels). A similar pattern of expression was also observed for *CTGF* and *ID3* (Fig. 3C, upper and lower panels). To determine whether these subgroups represented more ALK5- or ALK1- responsive cell populations, we evaluated the expression of ALK5- and ALK1 signature gene sets, chosen based on their strong dysregulation in our microarray analysis of HSVSMC and including well-established ALK5 & ALK1 target genes (signature gene sets listed in Supplemental Information, Table 4). UMAP visualization showed an opposing expression pattern for the two transcriptional signatures (Fig.3D and Fig.3F). Gene ontology analysis of the 100 most differentially expressed genes (top 30 most differentially-expressed genes shown in Supplemental Fig. 6) in the ALK5-dominant subgroup showed specific enrichment of terms associated with endoplasmic reticulum stress, chondroblast differentiation, regulation of smooth muscle cell migration & proliferation (Supplemental Fig.7A, FDR p<0.05). Interestingly, ‘negative regulation of wound healing’ was identified as an enriched term, in line with our observation of enhanced HSVSMC migration following pharmacological inhibition of ALK5 (Fig.2F). Conversely, the ALK1-dominant group (subgroup 0) showed enrichment of terms associated with senescence, positive regulation of cell aging and vasculogenesis (Supplemental Fig.7B, FDR p<0.05). Next, we used the scvelo RNA velocity toolkit to analyze the dynamic change in the transcriptional state of TGFβ-treated HSVSMC. Scvelo is a toolkit that can be used to model transcript dynamics in scRNA-seq datasets, by statistical modeling of RNA processing and decay using nascent transcript information present in the scRNA-seq data. Interestingly, RNA velocity analysis of TGFβ-treated HSVSMCs suggested that treated cells were differentiating along separate lineages, with the ALK5-dominant and ALK1-dominant subgroups at opposite poles of the low-dimensional embedding produced with UMAP as shown in Fig.4B (RNA velocity plot for untreated cells shown in Fig.4A).

**Figure 3.**
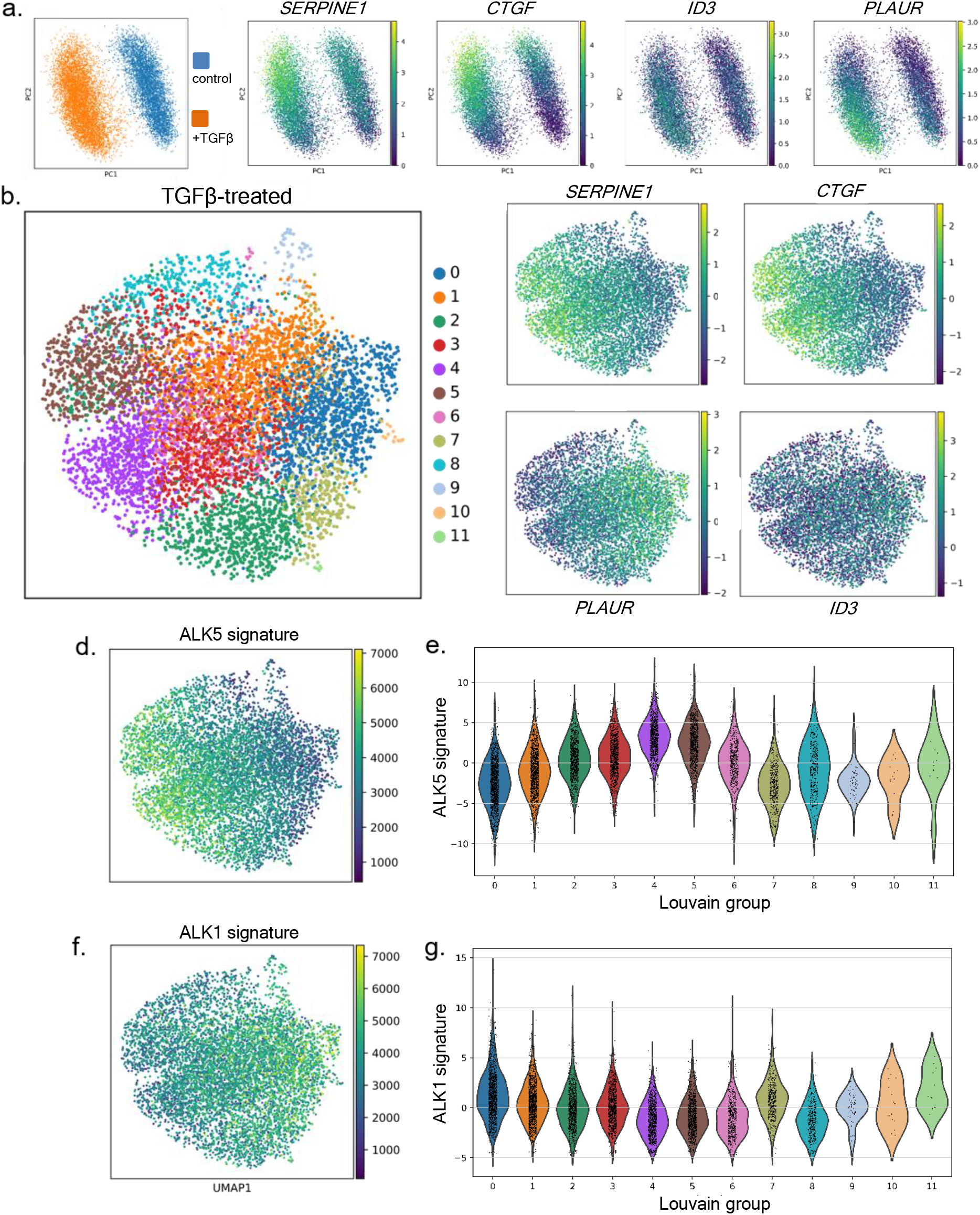
scRNA-seq analysis identifies nonoverlapping ALK5- and ALK1-dominant HSVSMC subgroups. Quiesced HSVSMC were stimulated with TGFβ for 24h prior to harvesting for scRNA-seq using the droplet-based 10x Genomics system, at an average read depth of ~50,000/cell. 5,245 untreated cells and 7,073 TGFβ-treated cells from the same patient were analysed using a Python pipeline (**A)** Cells were separated into two batches for analysis (TGFβ-treated/orange and untreated control/blue), with expression of SERPINE1, CTGF, ID3 and PLAUR mapped (**L-R**). **(B)** Graph-based UMAP clustering of TGFβ-treated cells using the Louvain method. (**C)** Expression of SERPINE1, CTGF, ID3 and PLAUR is mapped to the Louvain UMAP graph (upper and lower panels, gene names indicated above the plots). **(D)** Expression of an ALK5 signature gene set, mapped to the Louvain UMAP graph. Violin plots **(E)** indicate the expression of ALK5 signature genes across clusters. **(F)** Expression of an ALK1 signature gene set, mapped to the Louvain UMAP graph. Violin plots **(G)** indicate the expression of ALK1 signature genes across clusters.

**Figure 4.**
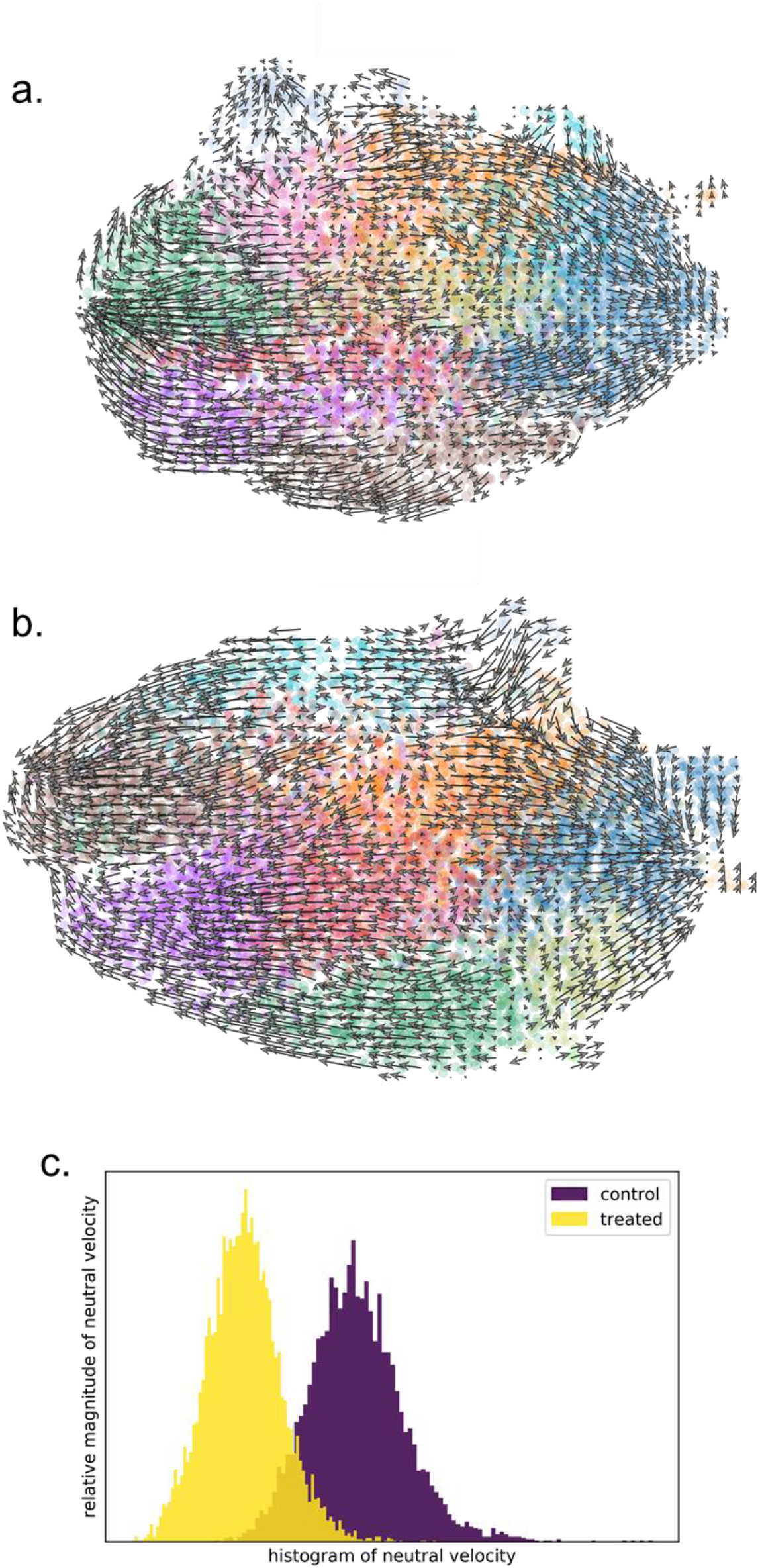
TGFβ treatment alters the transcriptional dynamics of HSVSMC. Quiesced HSVSMC were stimulated with TGFβ for 24h prior to harvesting for scRNA-seq using the droplet-based 10x Genomics system, at an average read depth of ~50,000/cell. RNA velocity analyses were performed on 5,245 untreated cells (**A**) and 7,073 TGFβ-treated cells (**B**) from the same patient. Arrows indicate directionality in the vector space, underlying colours correspond to Louvain grouping. Linear algebra was used to decompose velocity vectors for each cell into two components, a neutral velocity component and a directional component. Neutral directional components of treated and untreated cells were plotted against the relative magnitude of neutral velocity; increased neutral velocity corresponds to reduced directional movement in the RNA velocity analysis (**C**)

Having observed differences in the directionality of RNA velocity vectors between opposite poles of the UMAP plot for TGFβ-treated cells (Fig.4B), we next aimed to analyze how consistent the directionality of velocity vectors was, not only in the two-dimensional embedding visualized with UMAP plots, but also in the original velocity space. To this end, we devised a simple method based on linear algebra, whereby we decompose the velocity vector for each cell into two components: the directional velocity component (measured as flowing away from the cluster centre) and the neutral velocity component (measured as the velocity component minus directional velocity component). We observed that TGFβ-treated HSVSMC displayed lower neutral velocity than control, untreated HSVSMC, suggestive of higher directional velocity and, by extension, increased differentiation (Fig.4C).

### Genetic ablation or pharmacological inhibition of pSmad1/5 attenuates neointima formation in vivo

Previous studies in animal models of vascular injury have convincingly shown that ALK5-mediated TGFβ signaling promotes endothelial to mesenchymal transition [16], myofibroblast conversion [17] and extracellular matrix deposition [18]. However, no studies to date have addressed the role of ALK1 signaling during neointima formation. First, to identify whether ALK1-mediated TGFβ signaling in SMC might contribute to neointima formation *in vivo*, we used immunostaining to evaluate the localization of TGFβ signaling pathway components within SMC in an inter-positional cuff mouse model of vein graft disease [19]. Confocal microscopy revealed that TGFβ1, ALK5 and ALK1 were expressed in αSMA+ intimal SMCs 28 days following murine vein graft implantation (Fig.5A). Interestingly, although nuclear localization of both pSmad2/3 and pSmad1/5 was observed in αSMA+ intimal SMCs in remodelled vein graft tissue at the 28-day timepoint, only pSmad2/3 nuclear localization was present in native, un-grafted vena cava (Fig.5B). Vein graft neointima formation is initiated by the acute, reperfusion injury-mediated activation of the graft endothelium, which is followed by a period of SMC and extracellular matrix expansion [5]. As the contribution of different cell types to neointima formation varies widely depending on the experimental model used [20] and no single model can fully recapitulate human vein graft disease, we chose to also evaluate two models of arterial injury. Similar to our findings in murine vein grafts, we observed TGFβ1, ALK5 and ALK1 in αSMA+ intimal SMCs in wire-injured or ligated mouse carotid arteries (Supplemental Fig.8A and 8C) with nuclear localization of pSmad1/5 within SMC only evident in injured vessels (Supplemental Fig.8B and 8D). These data confirm the expression of ALK1 in intimal SMC and indicate that Smad1/5 is consistently activated in SMC during neointima formation *in vivo*.

**Figure 5.**
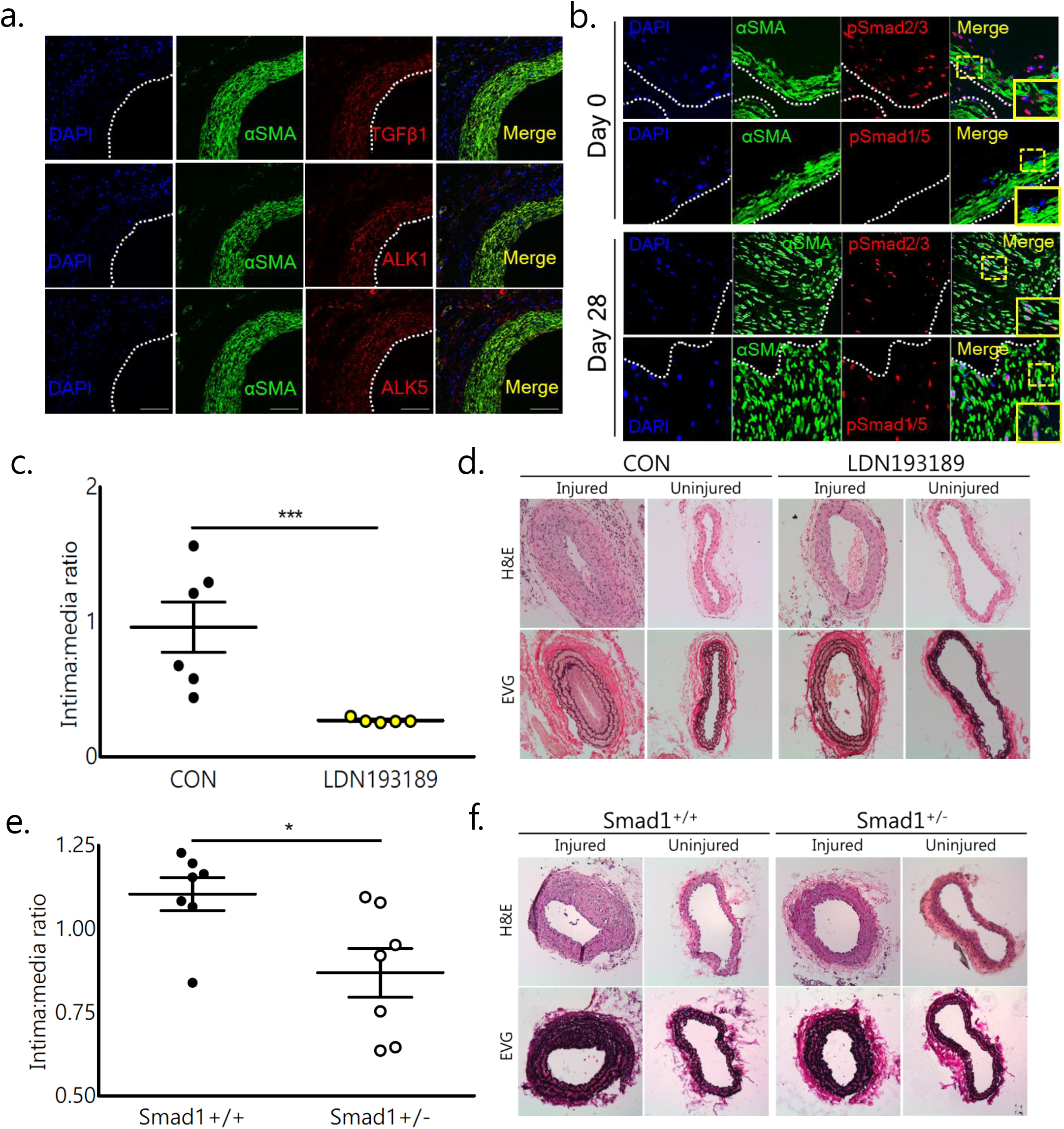
Smad1/5 activation promotes neointima formation following acute vascular injury in mice. **(A)** Immunostaining for TGFβ1, ALK1 or ALK5 in in remodelled mouse vein graft sections. (**B)** Dual immunostaining for pSmad2/3 and pSmad1/5 in pre-implantation (day 0, upper panels) and remodelled (day 28, lower panels) mouse vein grafts; SMC are identified using the SMC marker αSMA. Confocal images of immunostained sections are representative images from n=3 animals. Dotted line indicates location of the lumen. **(C, D)** Morphometric assessment of intima:media area ratio in LDN193189-treated (10mg/kg IP every 48h) or saline-treated control mice following acute vascular injury (n=5/6 per group). (**E/F**) morphometric assessment of intima:media area ratio following acute vascular injury in male Smad1^+/−^ mice or littermate controls (Smad^+/+^; n=8/group) *=p<0.05, ***=p<0.001 compared to wild-type/control by students T-test; IMR data represent mean ± SEM.

We next aimed to assess the contribution of pSmad1/5 signaling in SMCs to neointima formation by inducing acute vascular injury in male c57bl6/J mice treated with 10mg^kg-1^ LDN193189, which inhibits Smad1/5 phosphorylation downstream of ALK1/ALK2(IC_50_ ALK1 = 0.8nM, IC_50_ ALK2 = 0.8nM; [21, 22]) and Smad1^+/−^ mice or wild-type littermate controls. A substantial neointima was observed in injured carotid arteries from saline-treated c57bl6/J mice and Smad1^+/+^ mice (Fig.3D & 3F; 2.02±0.5×10^4^ and 1.8±0.2×10^4^ µm^2^, respectively, Supplemental Fig.9A & 9B). However, neointima formation was substantially reduced in LDN193189-treated mice compared to saline-treated controls (mean intima:media ratio/IMR = 0.26±0.01 vs 0.96±0.18, Fig.5C; p<0.001) and in Smad1^+/−^ mice compared to wild-type siblings (mean IMR = 0.87±0.07 vs 1.1±0.05, Fig.5E; p<0.05), indicating that activation of Smad1/5 following acute vascular injury promotes intimal hyperplasia.

### ALK1 and associated TGFβ signaling components are widely expressed in hyperplastic pre-implantation saphenous vein grafts

Having identified a pathological, pro-migratory role for ALK1/Smad1/5 through our *in vitro* and *in vivo* studies, we next sought to evaluate the expression of ALK1 and other components of the TGFβ signaling pathway in intact pre-implantation saphenous vein (SV) segments from CABG patients. Surprisingly, morphometric analyses revealed the presence of pre-existing intimal thickening (defined as an IMR >0.15) in over 88% of SV segments analyzed (Fig.6A & 6B; n=75, IMR range 0.067-1.37). The pathogenesis of vein graft IH is associated with the acquisition of a proliferative, synthetic phenotype by medial SMC, characterised by the downregulation of mature, contractile SMC proteins such as CNN1 and SMMHC [23]. Accordingly, confocal microscopy of immunostained SV sections with elevated IMR (>0.6) showed decreased CNN and SMMHC (Fig.6C; data not shown for SMMHC) but increased PCNA (proliferating cell nuclear antigen; Fig.8D) in the SV neointima compared to the media (n=8-10). qRT-PCR evaluation of laser-capture micro-dissected segments of SV neointima and media provided quantitative confirmation that genes associated with SMC contractility were downregulated in intimal segments, while genes associated with cell cycle progression were concordantly upregulated (Supplemental Fig.10B, n=8). Together, these confirm the presence of IH in pre-implantation SV segments from our cohort of CABG patients.

**Figure 6.**
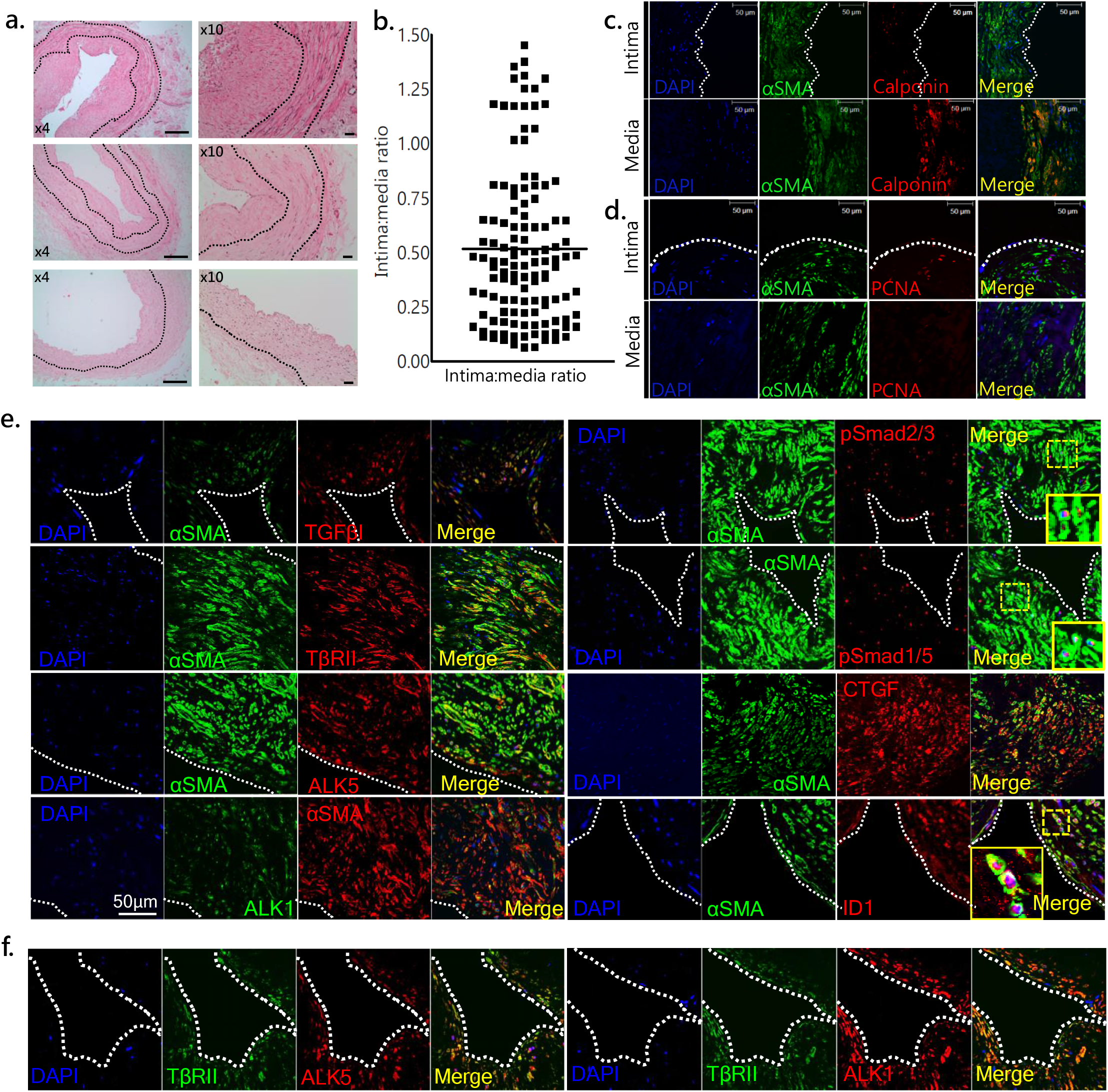
Histological analyses of saphenous vein grafts (SVG) from CABG patients show widespread pre-existing intimal hyperplasia and expression of TGFβ signaling pathway components. Surplus SVG tissue was obtained from a cohort of CABG patients (n=75). Intima and media areas and ratios (IMR) were calculated by performing morphometric analyses on H&E stained SVG sections (**B**) (representative H&E images from high[top]/medium[middle]/low[bottom] IMR SVG sections are shown in **A)**. Data are expressed as mean ± SEM. Immunostaining for the mature SMC marker calponin (CNN; **C**) and PCNA (**D**) in SVG sections with IMR >0.6 **(E)** Immunostaining for TGFβ1, TβRII, ALK5, ALK1, pSmad2/3, pSmad1/5, CTGF and ID1(target genes of ALK5 and ALK1, respectively), as well as colocalisation of TβRII and ALK5 (**F**, left panel) and ALK1 (**F**, right panel) in SVG sections with IMR>0.6. αSMA was used as a global marker for SMCs. Inset images show nuclear localisation of pSmads in αSMA+ SMCs. n=7-15 patient samples analysed, representative confocal microscopy images shown. Scale bars are indicated.

Finally, to identify whether ALK1 signaling in SMC might contribute to vein graft IH we evaluated the localization of TGFβ signaling pathway components within SV samples showing substantial pre-existing neointima formation (IMR>0.6; n=7-15). Immunostaining revealed that TGFβ1, TβR2, ALK5 and ALK1 were all highly expressed in αSMA+ intimal SMCs (Fig.6E), while dual staining for TβR2 and ALK5 or ALK1 showed strong colocalization between ALK1/ALK5 and TβR2(Fig.6F). Both pSmad2/3 and pSmad1/5 were localized to the nuclei of intimal SMCs, which also showed elevated expression of the transcriptional targets *CTGF* and *ID1* (Fig.6E). Furthermore, gene expression analysis of laser-capture microdissected SV segments confirmed the expression of the top transcriptional targets identified in our HSVSMC microarray analyses (*CTGF*, *FGF2*, *SCARA3* and *FADS1*; Supplemental Fig.11). Interestingly, we also observed abundant nuclear-localized pSmad2/3 and pSmad1/5 in αSMA+ intimal SMCs in three failed SVG specimens obtained from patients undergoing cardiac transplantation (Supplemental Fig.12A & 12B). Together, these data indicate that ALK1 expression in SMC within pre-implantation SV tissue from CABG patients may be clinically relevant to the pathogenesis of vein graft neointima formation and, ultimately, failure.

## Discussion

Although previous studies have extensively characterized the pro-fibrotic role of ALK5 in animal models of vascular injury [16–18], no studies to date have addressed the role of ALK1 signaling during neointima formation following CABG surgery. We report for the first time that ALK1 and its co-receptor endoglin, which can potentiate ALK1 signaling, are highly expressed in primary human vein graft SMCs from CABG patients (HSVSMC); we also show that TGFβ can bind both these receptors at the cell surface. Importantly, using ALK inhibitors and siRNA approaches we demonstrate that induction of Smad1/5 phosphorylation by TGFβ in HSVSMC is mediated by ALK1, with canonical Smad2/3 phosphorylation occurring downstream of ALK5. The ability of TGFβ to signal via two different type I receptors was first identified in ECs, with early studies showing that activation of ALK1/pSmad1/5 by TGFβ induced EC proliferation and migration whereas activation of ALK5/pSmad2/3 antagonized these processes [28]. While our results contrast with previous studies in cultured human aortic SMCs, where phosphorylation of Smad1/5 was shown to be primarily mediated by ALK5 [29], another group has recently demonstrated that TGFβ- induced Smad1/5 phosphorylation in new-born murine pulmonary artery SMCs involves ALK1 [30], suggesting that the ability of TGFβ to activate this signaling mechanism varies between SMC sub-types. This context-specificity is an established hallmark of TGFβ superfamily signaling and critically influences cellular responses to TGFβ; moreover, studies show that type I receptors bind different ligands in different contexts, with ALK1 able to act downstream of bone morphogenetic protein-9 (BMP9) and BMP10 as well as TGFβ [31, 32]. Interestingly, *Daly et al* [33] reported that heterotetrameric receptor complexes incorporating ALK5 and ALK1 generate mixed pSmad complexes which have lower affinity for BMP response elements in canonical Smad1/5 target genes such as *ID1* and *ID3*. This has raised questions around the functional relevance of ALK1:TGFβ signaling. Whilst we cannot exclude the generation of mixed Smad complexes in our experiments, the nuclear localization of pSmad1/5 and sustained induction of *ID1* expression (and *ID3*; data not shown) following TGFβ stimulation clearly indicates that activation of ALK1 is functionally significant in HSVSMC.

Our qRT-PCR validated, whole genome expression profiling of HSVSMC provides further evidence that ALK1 and ALK5 are both key mediators of TGFβ signaling in vein graft SMCs, showing that TGFβ regulates distinct transcriptional pathways by signaling via these two receptors. Specifically, we observed that the upregulation of SMC contractility genes by TGFβ was driven by ALK5, associated with enhanced pS20-MLC expression and corroborated using an established SMC contractile differentiation model. Interestingly, our scRNA-seq analyses suggested that treated cells were differentiating along separate lineages, with the ALK5-dominant and ALK1-dominant subgroup separated in the UMAP embedding space. Computational estimation of the directionality of RNA velocity vectors indicated reduced neutral velocity in TGFβ-treated HSVSMC compared to control, untreated HSVSMC, indicative of increased differentiation following TGFβ treatment. We would like to note however, that estimation of RNA velocity vectors is still an active area of research and our method of RNA vector decomposition would have to be tried on larger sample sizes before it can lead to any definitive biological conclusions.

Previous studies performed in SMCs of arterial origin have shown that activated Smad2/3 complexes interact with Smad binding elements in the αSMA (*ACTA2*?), SM22α (*TAGLN*) and SMMHC (*MYH11*) promoters [34]. Our findings therefore show that this transcriptional mechanism is highly conserved across venous and arterial beds, highlighting a fundamental role for ALK5 in the regulation of SMC contractility. We also found similarities between our ALK5 target gene dataset and those identified from previous microarray analyses performed in ECs, in which *CTGF* and collagen genes were shown to be specific targets of the ALK5 pathway. This suggests wider conservation of TGFβ:ALK5 gene targets across cell types [35, 36]. Conversely, the ALK1-specific transcriptional targets we identified showed little overlap with known TGFβ or ALK1 target genes [36, 37], and included genes such as *SCARA3,* which encodes a scavenger receptor that promotes the uptake of modified lipoproteins and plays an important role in fatty streak development during atherogenesis [38]. This may be linked to differences in experimental approach: previous studies overexpressed a constitutively active form of ALK1 in ECs, whilst we employed a recently-developed pharmacological ALK1 inhibitor (K02288) to pinpoint TGFβ-specific targets downstream of ALK1 in CABG patient SMC. Alternatively, this may indicate that the response to ALK1 activation by TGFβ is more sensitive to cell context, varying according to the presence or absence of transcriptional coregulators. Interestingly, the pro-migratory transcription factors Sox4 and Runx1 (Runt-related transcription factor 1) ([39][40]), which can act as DNA binding co-factors for Smad1/Smad5, are upregulated by TGFβ in HSVSMCs (2.1- and 3-fold, respectively; data not shown); by integrating TGFβ:ALK1 signaling, co-regulators such as Sox4 and Runx1 may help promote HSVSMC migration. In support of this hypothesis, our microarray analyses and wound healing assays identified a novel role for ALK1 in the regulation of HSVSMC migration. SMC migration is integral to vein graft neointima formation, during which proliferative medial SMCs move into the expanding intima. ALK1 expression and activation in HSVSMCs may therefore be a pathophysiological response to injury, promoting SMC migration and negative, inward vein graft remodeling.

In agreement with previous reports [10, 18] we found that TGFβ, ALK5 and ALK1 are expressed by neointimal SMCs in injured mouse carotid arteries and remodeled vein grafts. We also observed that pSmad2/3 and pSmad1/5 were localized to the nuclei of SMCs in mouse vein grafts, in correlation with the *Cooley et al* study on EndMT in VGF [41]. Interestingly, while ALK5 signaling was active in both non-injured and injured vessels, ALK1 signaling was only activated following vascular injury, evidenced by the absence of nuclear pSmad1/5 in uninjured vessels. Whilst BMPs can also activate Smad1/5 via ALK2, 3 and 6, previous studies indicate that only ALK1 expression is upregulated in wire-injured arteries 28 days following injury [10]. Together with the substantially reduced neointima formation we observed in mice treated with the ALK1/2 inhibitor LDN193189 and in heterozygous null *Smad1* mice, these data indicate that activation of the ALK1/Smad1/5 pathway is involved in the acute phase response to vascular injury. The identity of the ligand(s) triggering Smad1/5 activation *in vivo* remains to be elucidated; although our *in vitro* work clearly demonstrates that TGFβ is an active ligand for ALK1 in HSVSMC, ALK1 also binds BMP9 and BMP10 with high affinity [42]. The role of alternate ALK1 ligands during neointima formation following acute vascular injury therefore requires further investigation. However, we were unable to detect BMPRII expression in injured mouse arteries following carotid ligation (data not shown), which suggests a role for TGFβ:ALK1 in the phosphorylation of Smad1/5 in SMCs during neointima formation after vascular injury.

To evaluate the clinical relevance of ALK1 expression in cultured HSVSMC, we performed detailed histological assessments of surplus pre-implantation SVG tissues from a cohort of 75 patients undergoing CABG surgery, reasoning that the presence of ALK1 in SMC within these samples may provide further evidence supporting a role for this pathway in vein graft neointima formation. We were surprised to observe that the vast majority of these vessels showed significant pre-existing IH, with IMRs measuring >0.15 in 88% of the vein graft segments analyzed. Interestingly, several studies including *ex vivo* organ culture experiments [43] and ultrasonographic studies of CABG grafts *in situ* [44, 45] indicate that pre-existing intimal thickening within SVG can act as a substrate for accelerated neointima formation following vein graft implantation and increase the risk of failure. Follow-up studies of our cohort of CABG patients will be of value in determining the clinical impact of our histological observations. Of note, ALK1 was highly expressed in αSMA+ intimal SMCs from the most hyperplastic vein graft segments (IMR>0.6), associated with reduced mature SMC marker expression and increased markers of proliferation. Although we cannot exclude the contribution of alternate ALK1 ligands, clinical studies of re-stenotic arteries clearly show, that TGFβ is acutely and chronically upregulated at vascular injury sites [reviewed in Bradshaw et al, 2018]. Together with our data showing localization of ALK1, TβRII, pSmads and target genes to vein graft SMCs, this is strongly suggestive of a role for TGFβ:ALK1 in the development of vein graft IH.

In summary, the present study shows that TGFβ activates two distinct Smad-mediated signaling pathways downstream of ALK5 and ALK1 in vein graft SMC from CABG patients, inducing receptor-specific transcriptional networks associated with SMC contractility and migration, respectively. Smad1/5 activation is identified as a pathological hallmark of the SMC response to acute vascular injury, promoting neointima formation and inward remodeling. Tipping the balance of TGFβ signaling away from ALK1 may represent a novel therapeutic strategy for vein graft disease, resulting in a more favorable, contractile SMC phenotype during vein graft remodeling after CABG surgery.

### Methods

An expanded Methods section can be found in the supplemental materials.

## Author contributions

ACB, AHB, SAN, MJH and PTD designed the studies. ACB, ELL, JTS, DJK, ASS and MT performed the experiments, CD and SEF provided clinical material for the study and NM provided Smad1^+/−^ mice. ELL, JTS, MM, JmC and ACB analyzed the data. PH oversaw the 10X single cell assay and performed the initial CellRanger based analysis. AK and MP constructed the Python pipeline and performed bioinformatics analysis of scRNAseq data. ACB and ELL wrote the manuscript, which was edited by all authors.

## Acknowledgements

The authors thank Elaine Friel and Andrew Carswell for technical assistance and Prof. Kohei Miyazono (University of Tokyo) for whole antisera used in the radioligand binding assays. We are very grateful for the support provided by the NHS GG&C Biorepository, and acknowledge the fantastic generosity of the CABG patients who provided tissue samples for our study. The scRNAseq experiment was carried out at Glasgow Polyomics research facility at the University of Glasgow.

