## Supplementary material for "Transforming growth factor-beta signaling via ALK1 and ALK5 regulates distinct functional pathways in vein graft intimal hyperplasia": Materials and Methods

### Supplemental methods

**Human tissue collection.** Surplus pre-implantation saphenous veins from patients undergoing CABG surgery at the Golden Jubilee National Hospital in Glasgow were collected after ethical approval in collaboration with the NHS Greater Glasgow & Clyde Biorepository (REC reference 16/WS/0207; Bio-repository project reference 107). Immediately after removal, vein segments were stored in sterile saline and transported to the tissue culture laboratory. After removing excess fat and adventitia, sections of vein were either fixed in 4% paraformaldehyde (v/v in phosphate-buffered saline/PBS; Sigma Aldrich, Gillingham, UK) embedded in paraffin wax for immunohistochemical analyses or utilized for smooth muscle cell (SMC) purification.

Human vein grafts were freshly dissected from explanted hearts at the time of cardiac transplantation in Sheffield and frozen immediately or fixed in 4% w/v formaldehyde in PBS and embedded in paraffin wax (REC reference 12/NW/0036, North West Lancaster REC).

**Human saphenous vein smooth muscle cell (HSVSMC) purification and culture.** HSVSMC were isolated from human saphenous vein segments within 24 hours of surgery using the explant technique as described in [1]. HSVSMC were cultured in SMC growth medium (Smooth Muscle Cell Growth Medium 2 containing Smooth Muscle Cell Growth Medium 2 Supplement Mix, Promo Cell, Heidelberg, Germany) supplemented with 15 % (v/v) foetal bovine serum (FBS), 2 mM L-glutamine, 100 I.U/mL penicillin and 100 µg/mL streptomycin at 37 °C in a humidified atmosphere containing 5 % $CO_2$ to allow SMC outgrowth. To induce quiescence, HSVSMC were cultured for 72 hours in SMC low serum medium (Dulbecco's Modified Eagles Medium supplemented with 0.2% v/v FBS, 2 mM L-glutamine, 100 I.U/mL penicillin and 100 µg/mL streptomycin; ThermoFisher Scientific, Paisley, UK). Cells were utilized between passages 3 and 5 for all experiments.

**HSVSMC contractile differentiation model.** HSVSMC were cultured in 6- or 12-well tissue culture plates in SMC growth medium until they reached 90% confluency. SMC growth medium was then replaced by Media 231 (ThermoFisher Scientific) supplemented with smooth muscle differentiation supplement (SMDS; ThermoFisher Scientific), 100 I.U/mL penicillin and 100 µg/mL streptomycin, with a parallel 'proliferative SMC' control group remaining in 15% FCS SMC growth medium. RNA and protein were harvested at indicated time-points.

**HSVSMC stimulation.** HSVSMC were serum-starved for 72 hours in SMC low serum medium (see above). Cells were treated with 10 ng/ml of rTGFβ1 (stock in 4 mM HCl solution containing 1 mg/ml BSA; R&D Systems, MN, US). For ALK inhibitor studies, cells were subjected to 10 µM ALK1 small molecule inhibitor (K02288) or 10 µM ALK5 small molecule inhibitor (SB525334) unless otherwise indicated (10mM stock in DMSO; Tocris Bioscience, Bristol, UK) in the presence or absence of 10 ng/ml rTGFβ1. Unstimulated cells in 0.2% FBS alone (± DMSO for inhibitor studies) were used as a control.

**Transfection of HSVSMC with short interfering RNA (siRNA).** HSVSMC cultured in 12-well plates were reverse transfected with dicer-substrate siRNAs designed to target human ALK1 or ALK5 or a negative scrambled siRNA control (ALK1/ACVRL1: 5'GGACUAUAGACCACCCUUCUAUGAT3' and 3'CUCCUGAUAUCUGGUGGGAAGAUACUA 5', ref#142431676, ALK5/TGFBR1: 5'CUCAUAGAGAUUUCAAAUCAAAGAA3' and 2' ACGACUAUAGACCACCCUUCUAUGAT 3', ref#142431673) (Integrated DNA technologies, Iowa, USA; final concentration 25nM), using Lipofectamine 2000 transfection reagent and Opti-MEM reduced serum medium according to manufacturer's instructions (ThermoFisher Scientific). Antibiotic-free transfection media were replaced with complete SMC growth medium 6 hours post transfection and cells were left to recover overnight before serum starvation and/or stimulation.

**Scratch migration assay.** Cell migration was assessed using a scratch wound-healing assay. Briefly, a single vertical scratch was induced in the cell monolayer of each well using a sterile 200 µL pipette tip. Horizontal lines were drawn on the outside bottom surface of each well to act as a guide for measurements. Cells were stimulated with rTGFβ1 for 20 hours. Scratched cells in 0.2% FBS and 15% FBS were used as negative and positive controls respectively. Images of the scratch (3 images/well) were taken directly above the guide line using an EVOS XL microscope immediately following the scratch (0 hours) and 20 hours post scratch. Image analysis was performed using Image J software whereby the distance between the edges of the wound was measured at successive points across the scratch. Migration at 20 hours was expressed as a percentage of the original scratch width (0 hours).

**Quantitative real-time reverse transcription polymerase chain reaction (qRT-PCR).** Total RNA was extracted from HSVSMC using the miRNeasy mini kit (Qiagen, Manchester, UK) following manufacturer's instructions. For mRNA analysis, RNA was reverse transcribed using the Taqman® Reverse Transcription Kit (ThermoFisher Scientific) and random hexamer primers according to manufacturer's instructions. PCR reactions were performed using the Taqman® mRNA Assay Kit (ThermoFisher Scientific). Each reaction contained 1× TaqMan® Mastermix, 1× mRNA probe (as listed in Table 1) and 1.5-2µL RT product in a total volume of 10µL. Real-time PCR was performed at 95°C for 10 minutes, followed by 40 cycles of 95°C for 15 seconds and 60°C for 1 minute. All PCRs were performed in duplicate wells per plate. The threshold cycle (Ct) was used to determine the relative quantities of each mRNA. Measurements were normalized to the housekeeper *UBC* (ΔCt) and the inverse log of ΔΔCt gave the relative fold change (denoted as RQ).

**Western immunoblotting.** HSVSMCs were lyzed in RIPA buffer (150 mM NaCl, 1% NP-40, 0.5% sodium deoxycholate, 0.1% SDS, 50mM Tris pH 8) containing 1× Complete Protease Inhibitor Cocktail (Roche Diagnostics, West Sussex, UK) and Phosphatase Inhibitor Cocktail (Sigma Aldrich). Samples were equalized for protein content following quantitation of protein using a bicinchronic acid (BCA) assay (Pierce, Cramlington, UK) according to manufacturer's instructions and fractionated by SDS-PAGE on 10% (w/v) NuPAGE® Novex® Bis-Tris resolving gels (ThermoFisher Scientific). An Amersham ECL Rainbow Molecular Weight Marker (GE Healthcare Life Sciences) was run on each gel as a reference for protein size. Transfer of proteins from the gel onto nitrocellulose membrane (Amersham Biosciences, Buckinghamshire, UK) was achieved via electrophoretic elution for 1 hour 30 minutes at 4°C at 100V. After blocking in Sea Block buffer (ThermoFisher Scientific) membranes were immunoblotted with primary antibodies (as listed in Table 2) overnight at 4°C, followed by immunoblotting with IRDye-conjugated secondary antibodies (Li-cor Biosciences, Cambridge, UK). Membranes were scanned on an Odyssey® CLx imaging system and analyzed using Image Studio Lite Ver 4.0. Where indicated, bands were normalized to GAPDH loading control.

**Immunocytochemistry (ICC).** Fixed cells on chambered culture slides (Fisher Scientific, Loughborough, UK) were permeabilized with PBS containing 0.1% (v/v) (Sigma Aldrich) Triton X-100 (PBS-Triton) and then blocked in 10 % (v/v) goat serum (Vector Laboratories, Peterborough, UK)/PBS-Triton for 30 minutes at room temperature (RT). Following incubation, blocking solution was replaced with 1 % (v/v) goat serum/PBS-Triton containing specific primary antibodies (or corresponding IgG) and left to incubate overnight at 4 °C (antibodies listed in Table 3). Following incubation with fluorescently-conjugated secondary antibodies (AlexaFluor™, ThermoFisher Scientific) cells were mounted with glass coverslips using ProLong® Gold Antifade Mountant with DAPI (Molecular Probes, Oregon, USA). Cells were imaged on a Zeiss 510 LSM confocal microscope (Zeiss, Oberkochen, Germany). Settings on the microscope were kept constant for each related series of images, including the IgG control, to allow a fair comparison between relative fluorescence intensities.

**Radioligand binding and affinity crosslinking assay.** Iodination of TGFβ1 was performed using the chloramine T method and HSVSMC were subsequently affinity-labelled with the radioactive ligand as described in [2]. In brief, cells were incubated on ice with $I^{125}$-labelled TGFβ1 for 3 hours. After incubation, cells were washed with ice cold PBS and cross-linking was performed using 54 mM disuccinimidyl suberate (DSS) and 3 mM bis(sulfosuccinimidyl)suberate (BS3) (Pierce, Netherlands) for 15 minutes on ice. Next, cells were washed, scraped and lyzed in detachment buffer (10 mM Tris-HCL, 1 mM EDTA, 10% glycerol, 10 µL/mL PMSF, pH 7.4). Lysates were incubated with antibodies against TβRII, ALK1, ALK5, Betaglycan and Endoglin (whole antisera; a kind gift from K. Miyazono) overnight at 4°C and immune complexes were precipitated using protein A-Sepharose (Amersham Biosciences). Samples were washed, boiled in SDS sample buffer, and subjected to SDS-PAGE. Finally, gels were dried and scanned with the Storm imaging system (Amersham Biosciences).

**Microarray analysis.** Human whole-genome gene expression profiling was performed on RNA from quiesced HSVSMC stimulated with 10ng/mL rTGFβ1 in the presence or absence of 10µM SB525334 or KO2288 for 24 hours (n=3 patients), using the Human HT-12 v4 Expression BeadChip (Illumina, California, USA). RNA was extracted from cells using a miRNeasy mini kit (Qiagen) according to manufacturer's instructions, and quality controlled on a Agilent 2100 Bioanalyser. Whole genome expression profiling on supplied RNA was carried out by AROS Applied Biotechnology (Aarhus,

Denmark). Briefly, the Illumina® TotalPrep® RNA amplification kit was used to generate biotinylated amplified RNA for hybridization with the Illumina Human HT-12 v4 Expression BeadChip array. Following hybridization, the BeadChip array was scanned on the Illumina® BeadArray Reader and the raw intensity values were inputted into Illumina GenomeStudio® software for analysis. Quantile normalized and background subtracted probe level intensity values were exported from GenomeStudio® software into R (https://www.r-project.org/). To avoid negative values and to reduce the influence of probes with low signal intensities, a constant was added and the data was log2 transformed. After quality checking the data, statistical analysis was performed using the R package, limma (linear models for microarray data) based on the multi-level experiment approach. 4-way venn diagrams of limma significant probes were produced based on a 5 % false discovery rate (FDR) cut-off. For each probe, a log fold change, a FDR and an average expression were generated for upload to Ingenuity Pathway Analysis (IPA) where it was used for a functional analysis of differentially expressed probes. IPA software (Ingenuity Systems, www.ingenuity.com) was used to provide biological and functional information for probes/genes found to be differentially expressed following microarray data analysis. Microarray data have been deposited under accession number E-MTAB-7168 (ArrayExpress).
**Single-cell RNA sequencing.** HSVSMC were cultured in 6-well tissue culture plates in SMC growth medium until they reached 90% confluency. HSVSMC were serum-starved for 72 hours in culture media containing 0.2% FBS, then stimulated with rTGFβ1(10ng/ml) for 24 hours. Three wells of rTGFβ1-treated cells and three matched wells of untreated HSVSMC were harvested by trypsinisation, pooled, and re-suspended in serum-free penicillin/streptomycin-containing D-MEM media. Within 45 minutes of harvesting, resuspended cells were loaded into a Chromium Controller (10X Genomics) and cDNA was generated from polyadenylated RNA, then amplified and converted to Illumina compatible libraries according to standard protocol published in Single Cell 3’ Reagent Kits v2 User Guide (10X Genomics). These were then sequenced on an Illumina NextSeq500 with a sequencing depth of approximately 50,000 read pairs per cell. The read 1 and 2 were 26bp and 90bp long, respectively. Sequencing metrics, including depth and saturation, are shown in Supplemental Fig. X.
**Computational pipeline for scRNA-seq analysis.** FASTQs were processed using CellRanger (v2.1.1, Zheng et al, 2017 Nature Communications) with the hg38 genome assembly as the reference genome to generate a BAM file. The BAM file contains positioned-sorted, aligned reads, as well as error-corrected cellular and molecular barcodes. The aligner underlying CellRanger (v2.1.1) is the STAR aligner (Dobin et al, 2013 Bioinformatics). dropEst (Petukhov et al, 2018 Genome Biology) was then used to estimate molecular counts for each cell and to create the primary gene count matrix per sample. To perform RNA velocity analysis, dropEst was used to generate separate count matrices only containing UMIs classified as intronic, exonic or exon/intron spanning. The resulting R data frames were exported in MatrixMarket format. These matrices were loaded into our Python pipeline. Cellular barcodes not found in the equivalent primary count matrix generated by CellRanger were discarded and an aggregated AnnData matrix was generated to be used with ScanPy (Notebook 1, data import).

Detailed notebooks outlining the Scanpy pipeline and scRNA-seq bioinformatics analyses can be found on GitHub (https://nbviewer.jupyter.org/github/picrin/vascular_TGFB1/tree/master/analysis/). Cells were removed if fewer than 2000 genes were detected and genes were removed if detected in fewer than 100 cells. To remove poor-quality cells, defined as cells with high proportions of mitochondrial gene reads, ratios of mitochondrial genes to total gene counts were computed and cells with mitochondrial gene ratios >0.05 and/or total counts >100,000 were filtered out. Following filtering, the UMI counts were normalized to library size (total number of UMIs detected), scaled by 10,000 and log-transformed (Notebook 2, filtering). Highly-variable genes were then excluded by filtering based on dispersion, and variations in cell count as well as cell cycle confounders were regressed out using Scanpy's regress_out function (Notebook 3, regressing confounders). After separating the treated and untreated cell data into two samples, principal component analysis was performed on the Z scores of the normalized expression values within each sample. Graph-based UMAP clustering was performed using the top 40 principal components and 10 nearest neighbours. The Louvain method of clustering was used, as it effectively finds communities in large networks through modularity optimizations. ALK1, ALK5 and cVSMC were computed based on curated lists of known marker genes and transcriptional targets identified through our microarray analyses (signature genes listed in table 4 below). Using these gene lists, cells were ranked from the most to the least expressed, and the sum of these ranks computed for each cell. This approach gives equal weight to all genes and performs better in practice than log ranks or other attempted approaches. Scatterplots and violin plots were generated using the MatPlotLib function in the Scanpy package (Notebook 4, clustering). RNA velocity and partition-based graph abstraction were computed using the scVelo package (Notebook 5, RNA velocity). Lists of the 30 most differentially expressed genes for gene ontology (GO) analysis were computed using Mann-Whitney-Wilcoxon tests with Benjamini-Hochberg correction (Notebook 6, Gene Ontology) and subsequently analyzed using the Gene Ontology Resource GO Enrichment Analysis tool (geneontology.org; genes listed in Table 5 below; Fischer's exact test with a FDR cutoff of $p<0.05$). Heatmaps were generated using Seaborn (Notebook 7, Heatmaps). Finally, differentiation distance was computed by decomposing the RNA velocity vectors for each cell into two components: the directional velocity component (measured as flowing away from the cluster centre) and the neutral velocity component (measured as the velocity component minus directional velocity component) (Notebook 9, Glasgow Differentiation Factor).

**Mouse carotid wire injury and ligation injury models.** All animal experiments were approved by the UK Home Office (PPL60/4429) and the local ethics committee and were performed according to the UK Home Office guidelines. The carotid wire injury model was performed as described by Linder et al [3]. Briefly, blunt dissection was used to expose then liberate the left carotid artery from the surrounding tissue in isoflurane-anaesthetized mice (2%/$O_2$). A braided non-absorbable silk suture (6-0, Fine Science Tools) was ligated around the proximal branch of the exposed carotid. In animals undergoing carotid ligation injury, wounds were sutured and mice recovered at this point. In mice undergoing wire injury surgery, a temporary vessel clamp was then placed at the distal end of the ligated carotid artery to prevent blood flow before an incision was made in the stretch of vessel between clamp and suture. A rigid polyamide line (2-0, Ethilon, Johnson&Johnson, Belgium) was then inserted into the arterial lumen, advanced to the aortic arch and pulled back 3 times. Mice were sacrificed using a schedule 1 method 14 days (for carotid ligation studies) or 28 days (for wire injury studies) post-surgery. Contralateral carotid arteries were used as controls. Following sacrifice and exsanguination via saline perfusion, carotid arteries were collected for fixation in 4% (w/v) paraformaldehyde (Sigma-Aldrich, UK) and histological analysis. Morphometric analyses were performed on serial 5µm sections (from 5 cross section levels at 100µm intervals) stained with Verhoeff-van Gieson or haematoxylin & eosin (H&E). Images of 3-4 sections from each level were captured using an EVOS imaging system (ThermoFisher Scientific) and neointima formation was analyzed using Image-Pro Analyzer 7.0 in a blinded manner. Intima and media area were calculated using a free hand tool in ImageJ, from an average of 15-20 sections per animal.

**Immunohistochemistry.** Paraffin embedded 5 µm sections were de-waxed in histoclear before rehydration and antigen retrieval using heat and citrate buffer (0.01M, pH 6). Sections were blocked and permeabilized with 15% goat serum in TBS-Tween20 (0.1% v/v) or TBS-Triton X-100 (0.05% v/v) before immunostaining with primary antibodies (concentrations as indicated in Table 6). Corresponding immunoglobulins (IgGs) (Invitrogen, Paisley, UK) were used as a control. Primary antibodies were detected with appropriate fluorescently-conjugated secondary antibodies (AlexaFluor, ThermoFisher Scientific) and imaged on a Zeiss 510 LSM confocal microscope. Settings on the microscope were kept constant for each related series of images, including the IgG control, to allow a fair comparison between relative fluorescence intensities.

**Laser Capture Microdissection (LCM)** Glass membrane slides (Zeiss, Oberkochen, Germany) were baked for 4 hours at 180°C to remove RNases before being treated with 0.01% v/v poly-l-lysine (Sigma Aldrich) in nuclease-free water (ThermoFisher Scientific) for 2 minutes and subjected to 30 minutes of UV light. A microtome was used to cut 8µm transverse sections from paraffin-embedded tissue blocks which were transferred to a water bath containing autoclaved, 0.1% DEPC-treated water at 40°C. Sections were mounted onto the glass membrane slides, deparaffinized and rehydrated through an ethanol gradient before being stained with cresyl violet (0.01%, ThermoFisher Scientific), dehydrated through ethanol and dried. An AX10 laser capture microscope (Zeiss) was used to isolate intima and media segments from vessel sections which were stored in adhesive caps (Zeiss) to be used for RNA isolation. RNA was extracted from LCM sections using the RNeasy FFPE kit (Qiagen) as per manufacturer's instructions. RNA extracted from LCM sections was reverse transcribed to cDNA using a Superscript II first strand synthesis kit with random hexamers according to manufacturer's instructions (ThermoFisher Scientific).

**Statistical analysis** Data are presented as mean ± standard error of the mean (SEM). For *in vitro* experiments, where >3 patient cell replicates were analysed (in triplicate), repeated measures one-way

ANOVA with Tukey's post-hoc tests were performed to determine significance between groups whilst taking into account patient-specific variation. In experiments where >2 independent experimental groups were analysed, one-way ANOVA with Tukey's post-hoc tests were utilized. For experiments comparing 2 groups, student's T tests (or paired T tests where relevant) were employed. Significance was set at $p<0.05$. qRT-PCR analyses were performed on a $Log_2$-transformed scale, using the $\Delta\Delta Ct$ method. All statistical analyses were performed using GraphPad Prism v.4 (GraphPad Software, USA), apart from statistical analyses of microarray data, which were performed using the R package in limma based on the multi-level experiment approach as indicated above (5% FDR cut-off).

**Table 1. Taqman® Gene Expression Assays**

| Gene | Assay ID | RefSeq GeneID |
|---|---|---|
| ACTA2 | Hs00426835_g1 | NM_001141945.2 |
| ACTB | Hs01060665_g1 | NM_001101.3 |
| ACVRL1 | Hs00953798_m1 | NM_000020.2 |
| CCND1 | Hs00765553_m1 | NM_053056.2 |
| CNN1 | Hs00959434_m1 | NM_001299.5 |
| CTGF | Hs01026927_g1 | NM_001901.2 |
| ENG | Hs00923996_m1 | NM_000118.3 |
| FADS1 | Hs00203685_m1 | NM_013402.4 |
| FGF2 | Hs00266645_m1 | NM_002006.4 |
| H1F0 | Hs00271174_s1 | NM_005318.3 |
| ID1 | Hs03676575_s1 | NM_002165.3 |
| MALL | Hs01073822_m1 | NM_005434.4 |
| MYH11 | Hs00975796_m1 | NM_001040113.1 |
| PPP1R18 | Hs00292978_m1 | NM_001134870.1 |
| SCARA3 | Hs00939871_m1 | NM_016240.2 |
| SERPINE1 | Hs01126606_m1 | NM_000602.4 |
| TAGLN | Hs1038777_g1 | NM_001001522.1 |
| TGFBR1/ALK5 | Hs00610320_m1 | NM_001130916.1 |
| TGFBR2 | Hs00234253_m1 | NM_001024847.2 |
| TGFBR3 | Hs00234257_m1 | NM_001195683.1 |
| UBC | Hs01871556_s1 | NM_021009.5 |

**Table 2. Primary antibodies used for Western immunoblotting**

| Protein | Antibody Source | Molecular weight (kDa) | **Conc. (mg/mL)** | Working dilution | **Supplier, Cat#** |
|---|---|---|---|---|---|
| pSmad1/5 | Rabbit | 60 | 0.5 | 1:500 | CST, 9516 |
| pSmad2 | Rabbit | 60 | 0.5 | 1:500 | CST, 3108 |
| Smad1 | Rabbit | 60 | 0.5 | 1:1000 | CST, 6944 |
| Smad2 | Rabbit | 60 | 0.5 | 1:1000 | CST, 5339 |
| ACTA2 | Rabbit | 47 | 0.2 | 1:500 | Abcam, ab5694 |
| CNN1 | Rabbit | 34 | 0.063 | 1:500 | Abcam, ab46794 |
| SM22α | Rabbit | 23 | 1 | 1:1000 | Abcam, ab14106 |
| pS20-MLC | Rabbit | 20 | 1 | 1:1000 | Abcam, ab2480 |
| Total MLC | Rabbit | 20 | 0.5 | 1:1000 | Invitrogen, 720325 |
| GAPDH | Rabbit | 37 | 1 | 1:2000 | CST, 2118 |

**Table 3. Primary antibodies used in Immunocytochemistry**

| Protein | Antibody Source | Clonality | **Conc. (mg/mL)** | Working dilution | **Supplier, Cat#** |
|---|---|---|---|---|---|
| pSmad 1/5/8 | Rabbit | Polyclonal | 0.2 | 1:100 | Santa Cruz, sc-12353 |
| pSmad 2/3 | Rabbit | Polyclonal | 0.2 | 1:100 | Santa Cruz, sc-11769 |

**Table 4. Signature genes for scRNA-seq analysis**

| Signature | Gene |
|---|---|
| **ALK5** | TRIB3 |
| | CTGF |
| | FGF2 |
| | SERPINE1 |
| | NOX4 |
| | TGFBR1 |
| **ALK1** | SCARA3 |
| | CYB561 |
| | KDELR1 |
| | ID3 |
| | PLAUR |
| | ACVRL1 |
| **cVSMC** | ACTA2 |
| | TAGLN |
| | CNN1 |
| | MYOCDN |
| | SMTN |

**Table 5. Top 30 differentially expressed genes in ALK5 & ALK1-dominant subgroups**

| ALK5 | ALK1 |
|---|---|
| SERPINE1 | WWTR1 |
| FGF2 | PSD3 |
| CTGF | ANKH |
| CALD1 | SNX9 |
| IGFBP3 | SPOCK1 |
| DDAH1 | S100A13 |
| CCDC80 | PLAUR |
| TPM1 | MCC |
| CYR61 | JARID2 |
| DLC1 | CNIH3 |
| NEK7 | VCAN |
| PTX3 | SLC20A1 |
| EXT1 | SH3BGRL3 |
| PXDC1 | MMP14 |
| FRMD1 | SH3PDX2B |
| CRIM1 | APCDD1L |
| DMD | GPC6 |
| SYNE1 | TRPS1 |
| PRKG1 | ABHD2 |
| CTPS1 | LINC00511 |
| LOX | HMGA1 |
| THBS1 | SMIM3 |
| FSTL1 | FMNL3 |
| ITGBL1 | MGP |
| CDH2 | COL6A3 |
| AFAP1 | PRRX1 |
| PDE1C | HMGA2 |
| LM07 | ZEB1 |
| SEPT11 | THBD |
| FGF14 | ZNF521 |

**Table 6. Primary antibodies used in Immunohistochemistry**

| Protein | Source | Clonality | Conc. (mg/ml) | Working dilution | Supplier, Cat# |
|---|---|---|---|---|---|
| αSMA | Mouse | Monoclonal | 0.07 | 1:200 | Dako, M0851 |
| | Rabbit | Polyclonal | 0.2 | 1:200 | Abcam, ab5694 |
| vWF | Rabbit | Polyclonal | 0.3 | 1:100 | Dako, A0082 |
| CNN1 | Rabbit | Monoclonal | 0.063 | 1:50 | Abcam, ab46794 |
| PCNA | Rabbit | Polyclonal | 0.5 | 1:50 | Abcam, ab18197 |
| TGFβRII | Mouse | Monoclonal | 0.2 | 1:50 | Abcam, ab78419 |
| TGFβ1 | Rabbit | Polyclonal | 1 | 1:100 | Abcam, ab36495 |
| ALK1 | Rabbit | Polyclonal | 0.1 | 1:20 | Atlas, HPA007041 |
| ALK5 | Rabbit | Polyclonal | 0.2 | 1:20 | Santa Cruz, sc-398 |
| pSmad 1/5/8 | Rabbit | Polyclonal | 0.2 | 1:20 | Santa Cruz, sc-12353 |
| pSmad 2/3 | Rabbit | Polyclonal | 0.2 | 1:20 | Santa Cruz, sc-11769 |
