## Supplemental Figures for "Transforming growth factor-beta signaling via ALK1 and ALK5 regulates distinct functional pathways in vein graft intimal hyperplasia"

Supplemental Figure 1.

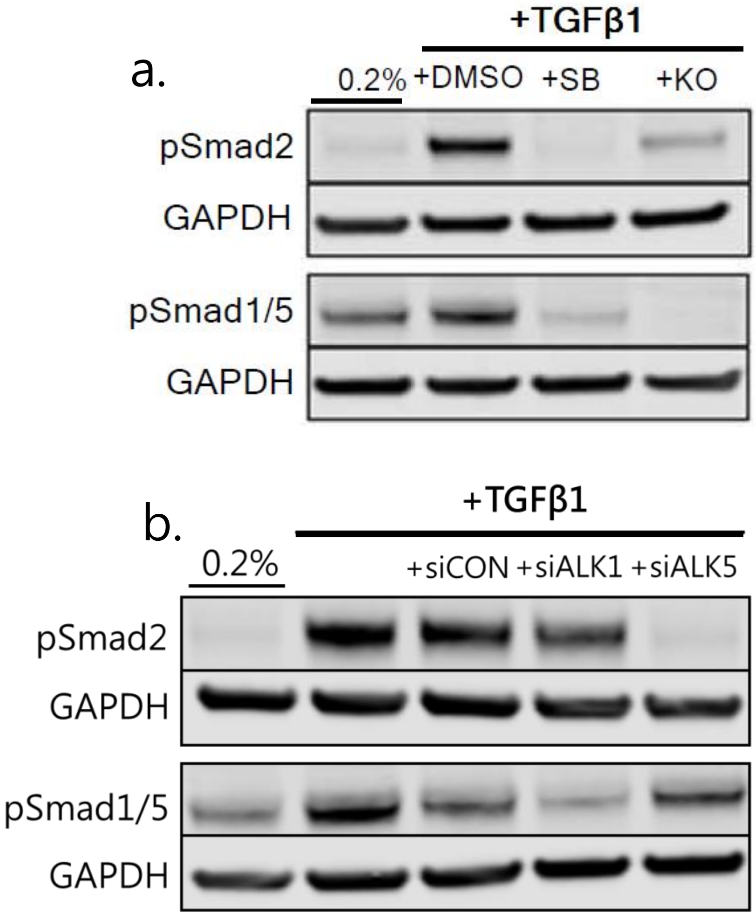


**Supplemental Figure 1. Effect of pharmacological inhibition or dsiRNA-mediated knockdown of ALK5/ALK1 on R-smad phosphorylation.** Serum-starved HSVSMCs from CABG patients were stimulated with TGFβ in the presence or absence of small molecule inhibitors of ALK5 (SB525334/SB) or ALK1 (K02288/KO) (**A;** 1h timepoint) or following dsiRNA-mediated knockdown of ALK5 or ALK1 (**B.** 1h timepoint) for western blot analysis of Smad2 or Smad1/5 phosphorylation. n=3; representative blots shown.

Supplemental Figure 2.

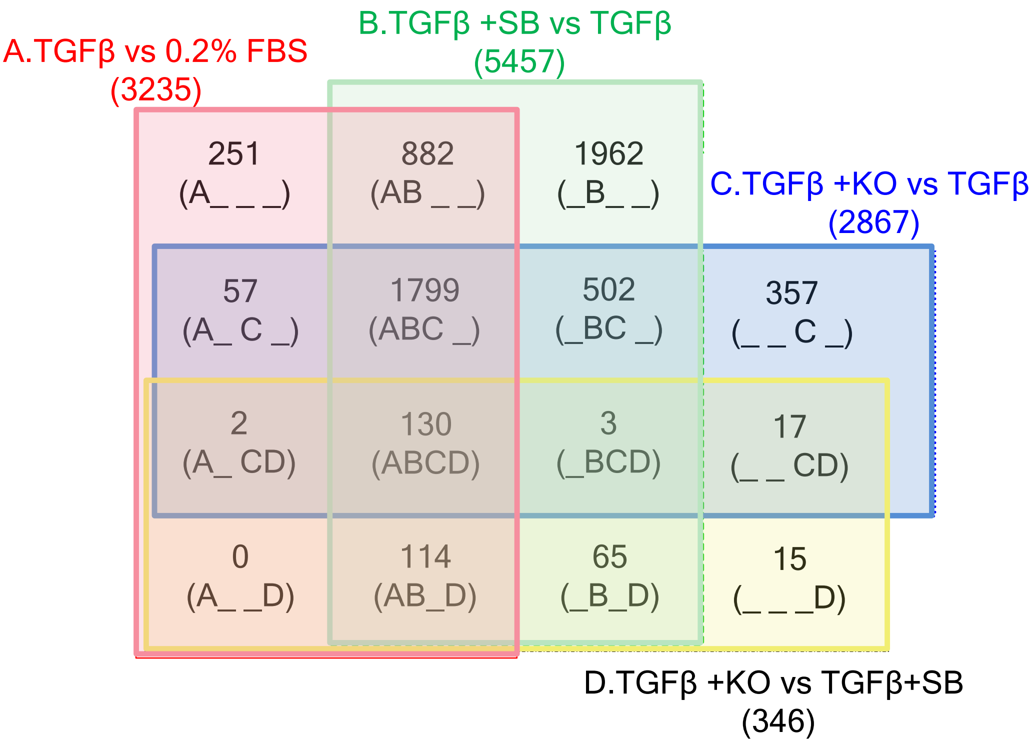


**Supplemental Figure 2. Four-way venn diagram of 5% FDR significant probesets from human whole genome expression profiling.** Serum-starved HSVSMCs from CABG patients were stimulated with TGFβ in the presence or absence of small molecule inhibitors of ALK5 (SB525334/SB) or ALK1 (K02288/KO) for 24h. Expression profiling was performed using the HT-12 v4 expression BeadChip (n=3 patients). Four-way venn diagrams of limma significant probes were produced in R based on a 5% FDR cut-off.

### Supplemental Figure 3.

A) Top downregulated genes following ALK5 inhibition (SB525334)

| Symbol | Gene Name | RefSeq GeneID | Fold Change |
|---|---|---|---|
| TRIB3 | Tribbles Pseudokinase 3 | NM_021158.3 | -5.62 |
| CTGF | Connective Tissue Growth Factor | NM_001901.1 | -4.76 |
| FGF2 | Fibroblast Growth Factor 2 | NM_002006.3 | -3.61 |
| MSC | Musculin | NM_005098.3 | -2.87 |
| COL16A1 | Collagen Type XVI Alpha 1 Chain | NM_001856.3 | -2.43 |
| SLC1A5 | Solute Carrier Family 1 Member 5 | NM_005628.1 | -2.31 |

B) Top upregulated genes following ALK5 inhibition (SB525334)

| Symbol | Gene Name | RefSeq GeneID | Fold Change |
|---|---|---|---|
| MALL | Mal, T-Cell Differentiation Protein Like | NM_005434.3 | 3.63 |
| BDKRB1 | Bradykinin Receptor B1 | NM_000710.2 | 2.55 |
| NDRG4 | NDRG Family Member 4 | NM_022910.1 | 2.38 |
| BTN3A3 | Butyrophilin Subfamily 3 Member A3 | NM_197974.1 | 1.83 |
| LM02 | LIM domain only 2 | NM_005574.2 | 1.81 |
| LUM | Lumican | NM_002345.3 | 1.8 |

C) Top downregulated genes following ALK1 inhibition (KO2288)

| Symbol | Gene Name | RefSeq GeneID | Fold Change |
|---|---|---|---|
| SCARA3 | Scavenger Receptor Class A Member 3 | NM_016240.2 | -1.76 |
| CYB561 | Cytochrome B561 | NM_001017917.1 | -1.5 |
| KDELR1 | KDEL Endoplasmic Reticulum Protein Retention Receptor 1 | NM_006801.2 | -1.41 |
| PPP1R18 | Protein Phosphatase 1 Regulatory Subunit 18 | NM_133471.1 | -1.4 |
| RFNG | RFNG-O-Fucosylpeptide 3-Beta-N-Acetylglucosaminyltransferase | XM_001132711.1 | -1.36 |
| GUK1 | Guanylate Kinase 1 | NM_000858.4 | -1.35 |

D) Top upregulated genes following ALK1 inhibition (KO2288)

| Symbol | Gene Name | RefSeq GeneID | Fold Change |
|---|---|---|---|
| FADS1 | Fatty Acid Desaturase 1 | NM_013402.3 | 2.14 |
| IDI1 | Isopentenyl-Diphosphate Delta Isomerase 1 | NM_004508.2 | 1.68 |
| H1F0 | H1 Histone Family Member 0 | NM_005318.2 | 1.55 |
| LAGE3 | L Antigen Family Member 3 | NM_006014.3 | 1.43 |
| NUDCD2 | NudC Domain Containing 2 | NM_145266.4 | 1.41 |
| GTDC1 | Glycosyltransferase Like Domain Containing 1 | NM_024659.2 | 1.4 |

**Supplemental Figure 3. Ingenuity pathway analysis of ALK1 or ALK5 gene targets in HSVSMC, showing top gene targets of ALK1 (C,D) or ALK5 signalling (A, B).** Human whole-genome expression profiling using the Illumina HT-12 v4 Expression BeadChip was performed on mRNA extracted from serum-starved HSVSMC stimulated for 24h with rTGFβ1 ± ALK5 or ALK1 kinase inhibitors (SB525334 or K02288, respectively; n=3). Fold changes of top gene targets is shown, based on limma significant probes identified using a 5 % FDR cut-off.

Supplemental Figure 4.

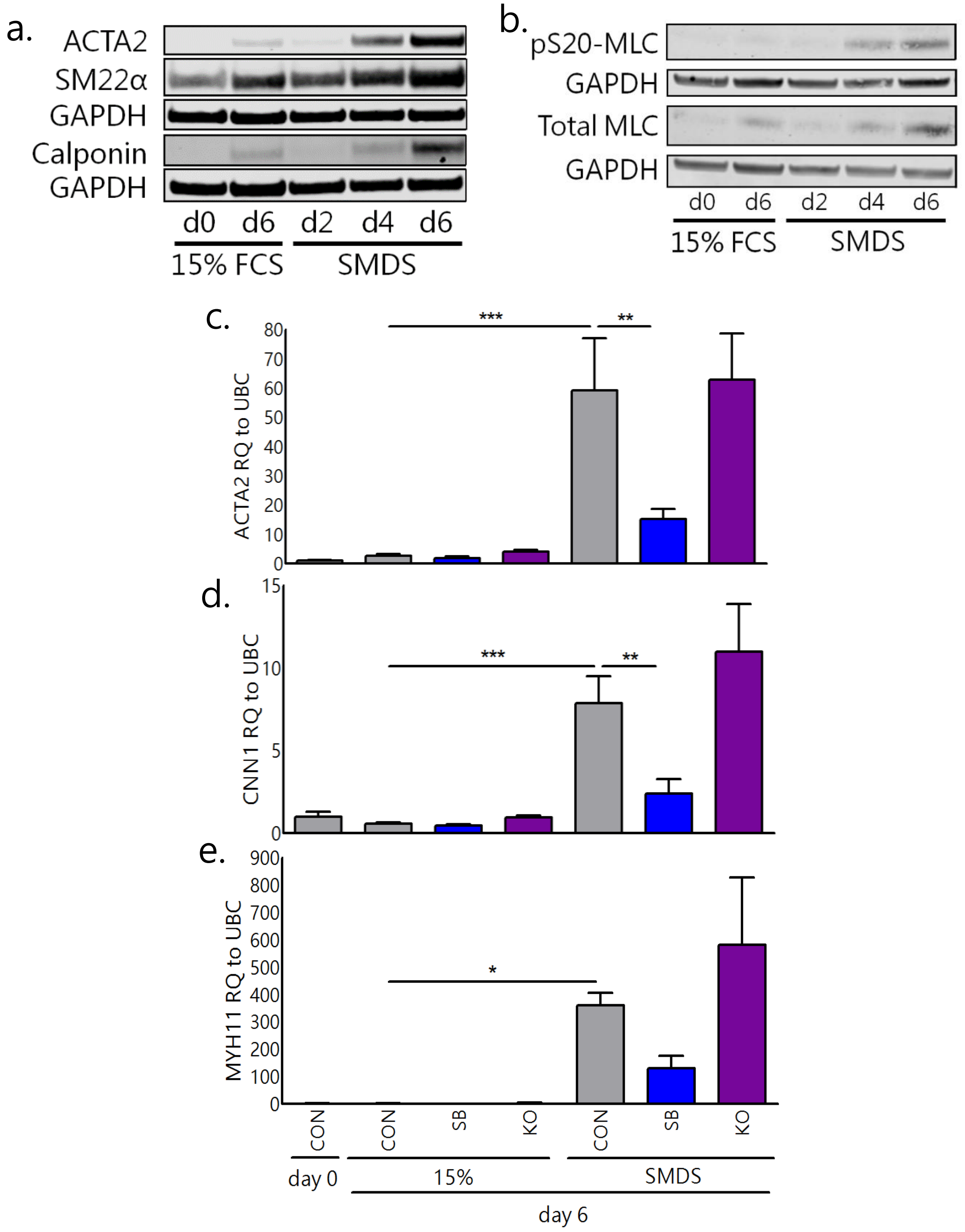


**Supplemental Figure 4. Effect of pharmacological ALK5/ALK1 inhibition on contractile differentiation of HSVSMCs.** HSVSMCs from CABG patients were incubated in 15% fetal calf serum (FCS) SMC growth medium or 1% FCS SMC differentiation medium (SMDS) for 6 days, in the presence or absence of small molecule inhibitors of ALK5 (SB/SB525334) or ALK1 (KO/KO2288). Contractile differentiation status was confirmed by immunoblot analysis of ACTA2 (α-smooth muscle actin), CNN1 (calponin) and SM22α (smooth muscle protein 22-α) (**A**) and phospho-myosin light chain (pS20-MLC)/total MLC (**B**). qRT-PCR analysis of ACTA2 (**C**), CNN1 (**D**) and MYH11 (**E**) in contractile (SMDS) or proliferative (15%) HSVSMCs treated with small molecule inhibitors of ALK5 (SB/SB525334) or ALK1 (KO/KO2288). Data in graphs represent mean ± SEM, analysed by repeated measures one-way ANOVA with Tukeys post-hoc tests; n=3 patients, *=p<0.05, **=p<0.01, ***=p<0.001.

### Supplemental Figure 5.

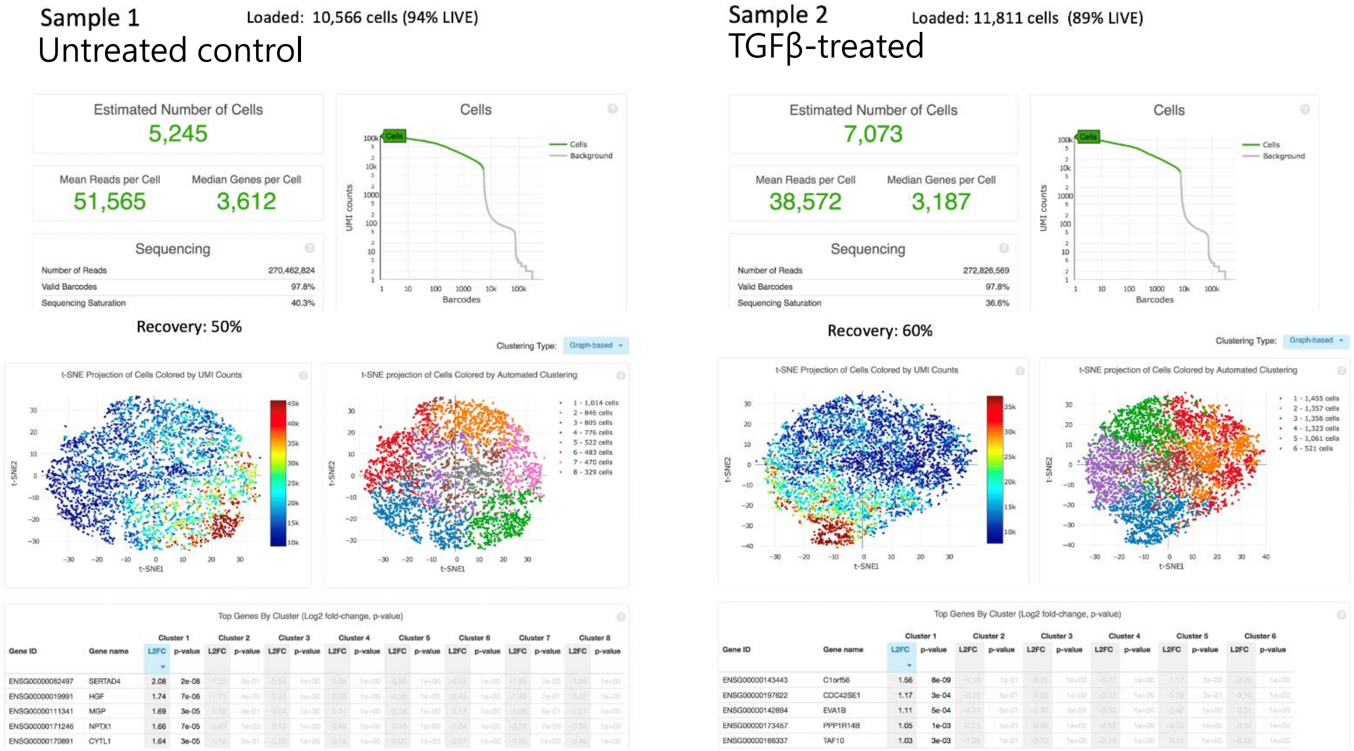


**Supplemental Figure 5. Quality control data from 10x genomics single cell RNA sequencing analysis of HSVSMC** Quiesced HSVSMC were stimulated with TGFβ for 24h prior to harvesting for scRNA-seq using the droplet-based 10x Genomics system, at a read depth of ~50,000/cell. 5,245 untreated cells (sample 1, L) and 7,073 TGFβ-treated cells (sample 2, R) were analysed. Plots show sequencing depth and coverage, with t-SNE projection of cells by UMI counts as analysed by the 10x Genomics CellRanger platform.

Supplemental Figure 6.

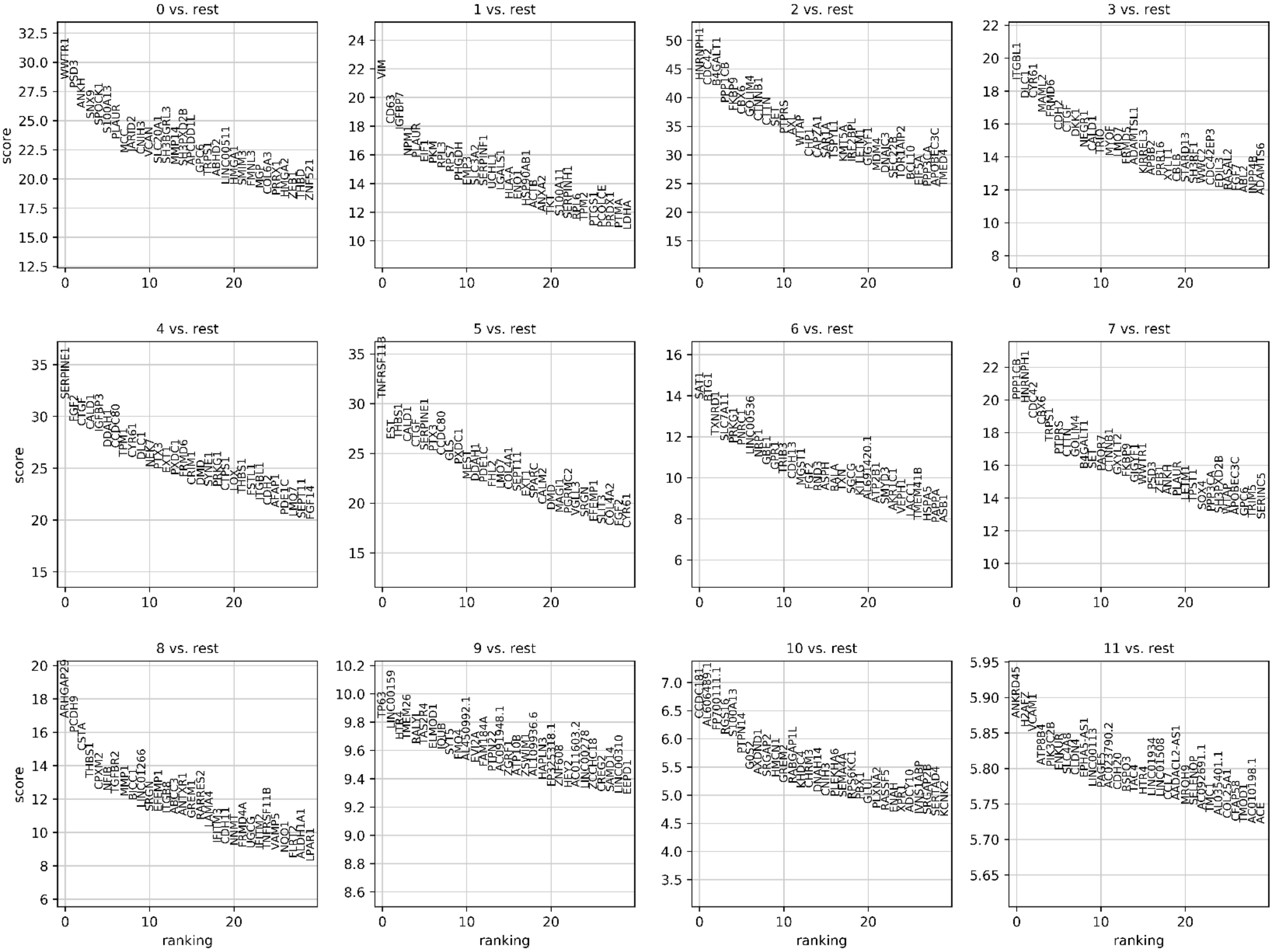


**Supplemental Figure 6. scRNAseq analysis identifies top 30 most differentially expressed genes in clusters of TGFβ-treated HSVSMC** Quiesced HSVSMC were stimulated with TGFβ for 24h prior to harvesting for scRNA-seq using the droplet-based 10x Genomics system, at a read depth of ~50,000/cell. UMAP clusters were identified through an unsupervised method using the Louvain algorithm. Individual plots list the 30 most differentially-expressed genes in each cluster.

### Supplemental Figure 7.

a.

| GO Rank | ALK5-dominant subgroup, GO biological process | No. genes | expected | Fold Enrichment | raw P value | FDR |
|---|---|---|---|---|---|---|
| 1 | response to methamphetamine hydrochloride | 2 | 0.01 | > 100 | 1.35E-04 | 0.0176 |
| 2 | glutamine catabolic process | 2 | 0.01 | > 100 | 2.25E-04 | 0.0248 |
| 3 | negative regulation of IRE1-mediated unfolded protein response | 2 | 0.01 | > 100 | 2.25E-04 | 0.0247 |
| 4 | chondroblast differentiation | 2 | 0.02 | 83.15 | 4.69E-04 | 0.0424 |
| 5 | negative regulation of endoplasmic reticulum unfolded protein response | 3 | 0.06 | 47.97 | 5.70E-05 | 0.00975 |
| 6 | negative regulation of smooth muscle cell migration | 4 | 0.11 | 37.8 | 6.82E-06 | 0.00205 |
| 7 | glutamine metabolic process | 4 | 0.11 | 36.15 | 7.98E-06 | 0.00231 |
| 8 | negative regulation of focal adhesion assembly | 3 | 0.09 | 34.65 | 1.33E-04 | 0.0176 |
| 9 | muscle cell cellular homeostasis | 3 | 0.1 | 29.7 | 2.00E-04 | 0.0229 |
| 10 | negative regulation of adherens junction organization | 3 | 0.1 | 29.7 | 2.00E-04 | 0.0228 |

b.

| GO Rank | ALK1-dominant subgroup, GO biological process | No. genes | expected | Fold Enrichment | raw P value | FDR |
|---|---|---|---|---|---|---|
| 1 | oncogene-induced cell senescence | | 0.01 | > 100 | 1.20E-04 | 3.34E-02 |
| 2 | senescence-associated heterochromatin focus assembly | | 0.01 | > 100 | 1.99E-04 | 4.52E-02 |
| 3 | positive regulation of cell aging | | 0.07 | 44.2 | 6.88E-05 | 2.28E-02 |
| 4 | positive regulation of collagen biosynthetic process | | 0.12 | 34 | 9.68E-06 | 4.53E-03 |
| 5 | positive regulation of collagen metabolic process | | 0.12 | 32.74 | 1.11E-05 | 5.03E-03 |
| 6 | regulation of collagen metabolic process | | 0.19 | 26.31 | 2.20E-06 | 1.30E-03 |
| 7 | response to fluid shear stress | | 0.16 | 25.26 | 2.82E-05 | 1.09E-02 |
| 8 | regulation of collagen biosynthetic process | | 0.17 | 23.89 | 3.44E-05 | 1.30E-02 |
| 9 | vasculogenesis | | 0.32 | 18.94 | 1.21E-06 | 8.00E-04 |
| 10 | regulation of phosphatidylinositol 3-kinase activity | | 0.25 | 15.79 | 1.55E-04 | 3.80E-02 |

**Supplemental Figure 7. Gene Ontology (GO) analysis of ALK5-dominant and ALK1-dominant scRNA-seq clusters of TGFβ-treated HSVSMC** GO enrichment analysis was performed using the top 100 most differentially expressed genes in the ALK5-dominant (subgroup 4) and ALK1-dominant (subgroup 0) clusters, using the GO Consortium gene ontology resource. A false discovery rate (FDR) rate cutoff of <0.05 was applied and results were ranked by Fold Enrichment score. The top 10 GO terms for ALK5-dominant (A) and ALK1-dominant (B) subgroups are presented.

### Supplemental Figure 8.

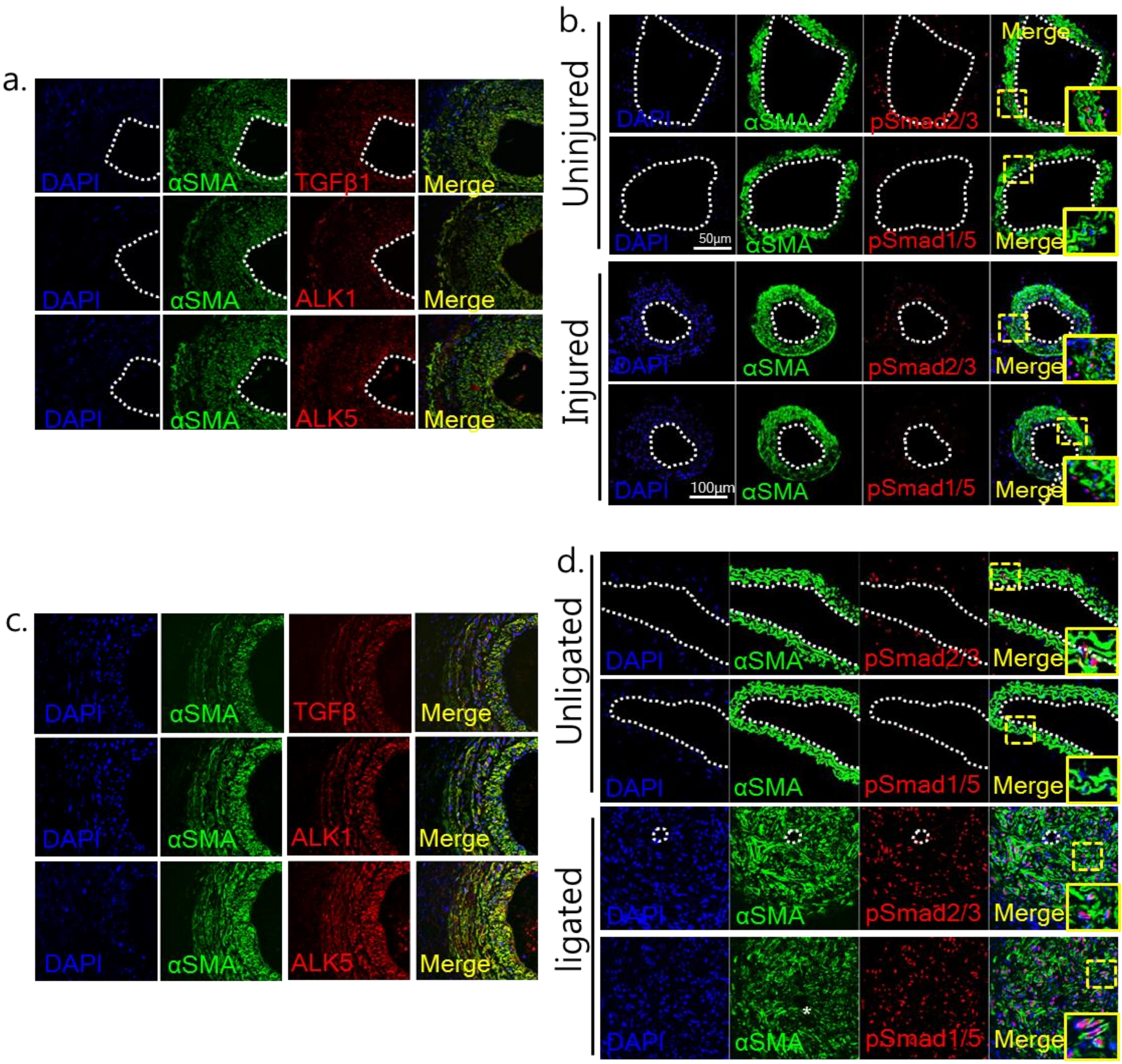


**Supplemental Figure 8. ALK pathway activation in mouse carotid wire injury (A,B) and ligation injury (C,D) models** Immunostaining for TGFβ1, ALK1 or ALK5 in αSMA+ SMC within wire-injured mouse carotid arteries **(A)** or ligated carotid arteries **(C)**. (**B/D**) dual immunostaining for pSmad2/3 and pSmad1/5 in uninjured mouse carotid arteries (**B**, **D**; upper panels) and wire-injured (**B**) or ligated (**D**) carotid arteries (lower panels); SMC are identified using the SMC marker αSMA. Confocal images of immunostained sections are representative images from n=4 animals/group. Dotted line indicates location of the lumen.

Supplemental Figure 9.

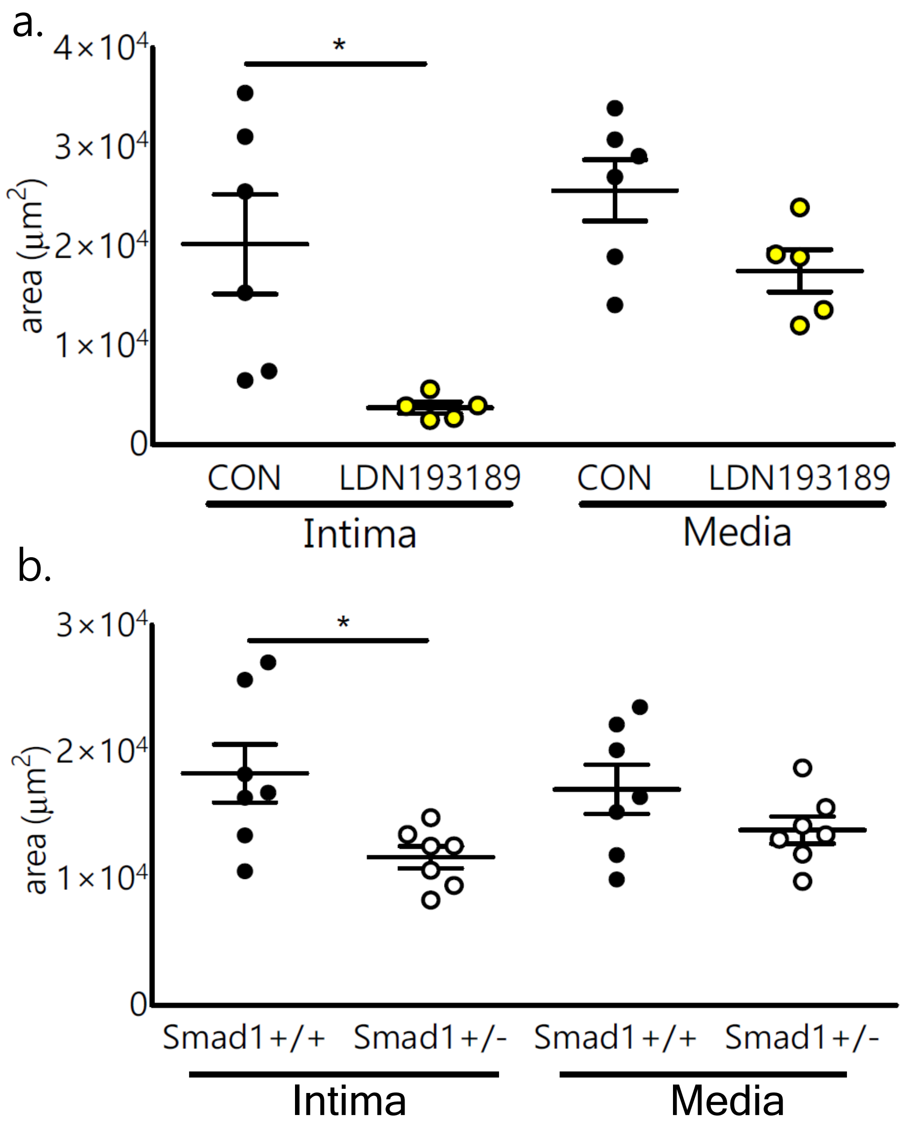


**Supplemental Figure 9. Morphometric analyses of carotid arteries from LDN193189-treated or Smad1$^{+/-}$ mice following acute vascular injury (A)** Intima and media area measurements from morphometric image analysis of carotid artery sections from LDN193189-treated mice (10mg$^{kg-1}$ IP) or saline-treated controls (n=5/6 per group). **(B)** Intima and media area measurements from morphometric image analysis of carotid artery sections from Smad1$^{+/-}$ mice and wild-type littermate controls (Smad$^{+/+}$; n=8/group). Data represent mean±SEM; *=p<0.05 compared to wild-type/control by 2-tailed students T-test.

Supplemental Figure 10.

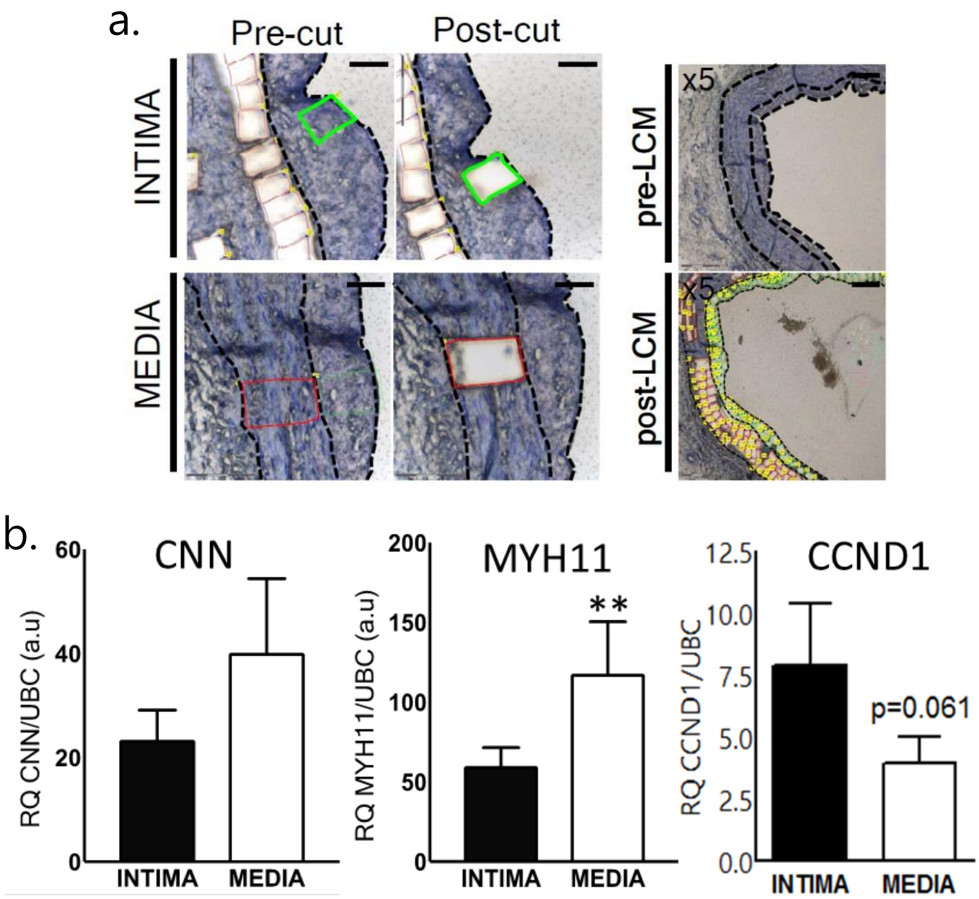


**Supplemental Figure 10. Gene expression of SMC or proliferation markers in intimal and medial segments from CABG SVG sections.** Measurement of calponin/*CNN*, smooth muscle myosin heavy chain (*MYH11*) and cyclin D1 (*CCND1*) mRNA expression by qRT-PCR (**B**; n=8) was performed on laser-capture microdissected samples of intimal and medial areas in SVG sections with IMR>0.6 (**A** shows areas of SVG intima or media excised using LCM). Data represent mean ± SEM; *=p<0.05 by paired t-test.

Supplemental Figure 11.

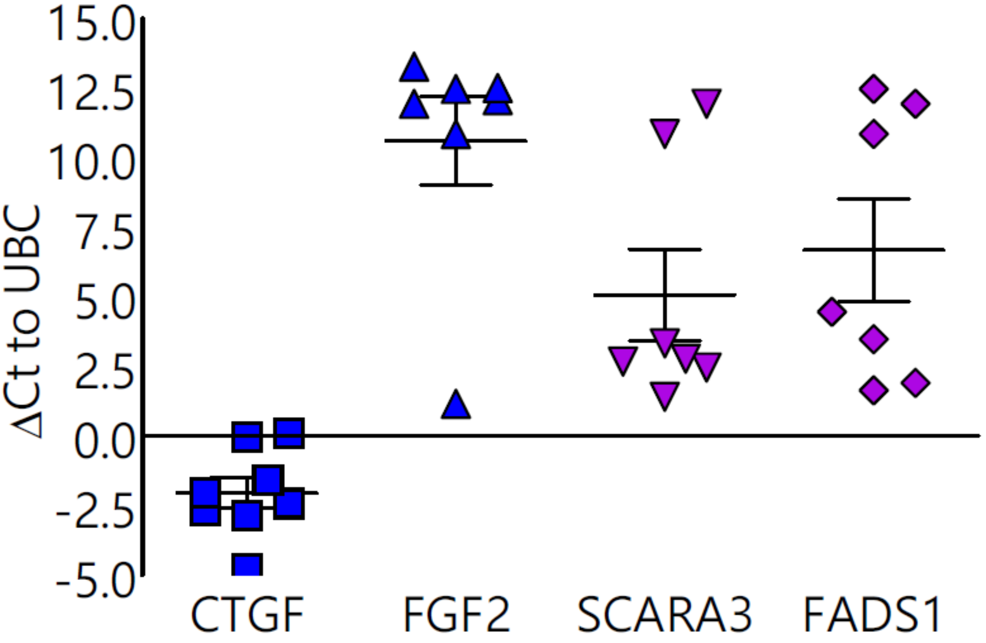


**Supplemental Figure 11. Gene expression of ALK5 (blue) or ALK1 transcriptional targets in CABG SVG sections.** mRNA expression of CTGF, FGF2, SCARA3 and FADS1 was performed by qRT-PCR on laser-capture microdissected samples of SVG from CABG patients. Bars show mean ΔCt relative to UBC control ± SEM (n=7-8).

Supplemental Figure 12.

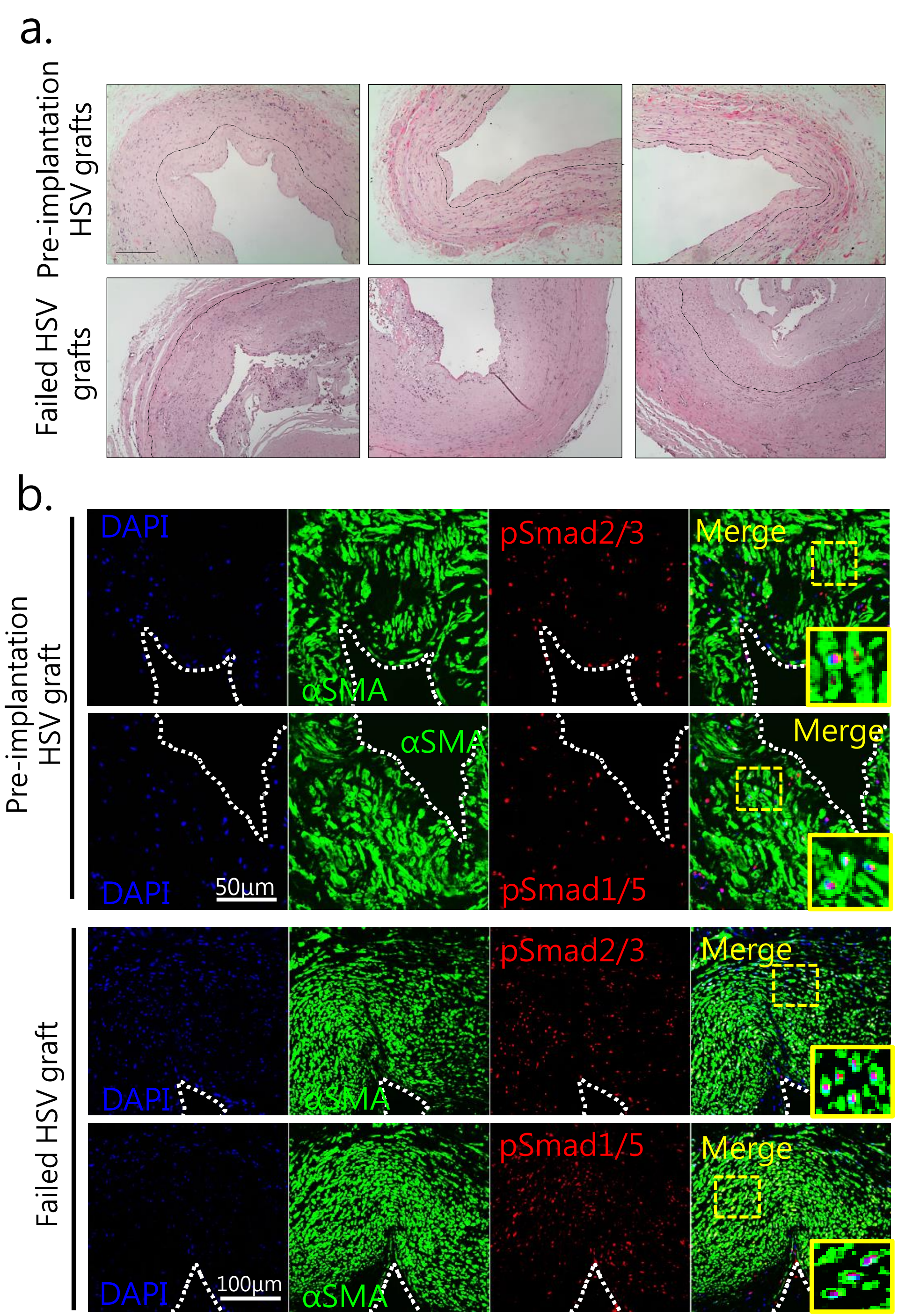


**Supplemental Figure 12. R-Smad phosphorylation in failed vein grafts from CABG patients.** Dual immunostaining for pSmad2/3, pSmad1/5 and the SMC marker αSMA in sections from pre-implantation SVG and failed SVG from CABG patients (**B**; H&E images in **A** show vessel morphology). Confocal images of immunostained sections are representative images from n=15 patients (pre-implantation) and n=3 patients (failed HSV grafts) respectively. Inset images show nuclear localisation of pSmads in αSMA$^{+}$ SMCs. Dotted line indicates location of the lumen. Scale bars are indicated.
